# Discovery of compounds inhibiting the ADP-ribosyltransferase activity of pertussis toxin

**DOI:** 10.1101/637801

**Authors:** Yashwanth Ashok, Moona Miettinen, Danilo Kimio Hirabae de Oliveira, Mahlet Z. Tamirat, Katja Näreoja, Avlokita Tiwari, Michael O. Hottiger, Mark S. Johnson, Lari Lehtiö, Arto T. Pulliainen

**Author notes:** CORRESPONDING AUTHOR INFORMATION Arto Pulliainen, Ph.D., Institute of Biomedicine, Research Center for Cancer, Infections, and Immunity, University of Turku, Kiinamyllynkatu 10, FI-20520, Turku, Finland, Fax: not available, Dr. Lari Lehtiö, Ph.D., Faculty of Biochemistry and Molecular Medicine & Biocenter Oulu, University of Oulu, Aapistie 7 B, FI-90220, Oulu, Finland, Fax: not available. Equal contribution (Yashwanth Ashok, Moona Miettinen).

## Abstract

Targeted pathogen-selective approach to antibiotic development holds promise to minimize collateral damage to the beneficial microbiome. The AB_5_-topology pertussis toxin (PtxS1-S5, 1:1:1:2:1) is a major virulence factor of *Bordetella pertussis*, the causative agent of the highly contagious respiratory disease whooping cough. Once internalized into the host cell, PtxS1 ADP-ribosylates α-subunits of the heterotrimeric Gαi-superfamily, thereby disrupting G-protein-coupled receptor signaling. Here, we report the discovery of the first small molecules inhibiting the ADP-ribosyltransferase activity of pertussis toxin. We developed protocols to purify mg-levels of truncated but highly active recombinant *B. pertussis* PtxS1 from *Escherichia coli* and an *in vitro* high throughput-compatible assay to quantify NAD^+^ consumption during PtxS1-catalyzed ADP-ribosylation of Gαi. Two inhibitory compounds (NSC228155 and NSC29193) with low micromolar IC_50_-values (3.0 µM and 6.8 µM) were identified in the *in vitro* NAD^+^ consumption assay that also were potent in an independent *in vitro* assay monitoring conjugation of ADP-ribose to Gαi. Docking and molecular dynamics simulations identified plausible binding poses of NSC228155 and in particular of NSC29193, most likely owing to the rigidity of the latter ligand, at the NAD^+^-binding pocket of PtxS1. NSC228155 inhibited the pertussis AB_5_ holotoxin-catalyzed ADP-ribosylation of Gαi in living human cells with a low micromolar IC_50_-value (2.4 µM). NSC228155 and NSC29193 might prove useful hit compounds in targeted *B. pertussis*-selective drug development.

## INTRODUCTION

Whooping cough, i.e. pertussis, is a globally distributed acute respiratory disease ^1^. Whooping cough affects all age groups ^1^. However, young children are the most affected age group where the disease may lead to death despite hospital intensive care and use of antibiotics ^1^. Worldwide estimates of cases and deaths in children younger than 5 years in 2014 were 24.1 million and 160.700, respectively ^2^. Despite efficient global vaccine campaign whooping cough remains endemic and recent data, e.g. from USA ^3^, indicate that the number of cases is increasing. Moreover, macrolide resistant *B. pertussis* strains have been reported ^4, 5^. Sizeable outbreaks have also been witnessed ^1^, but the reasons for the resurgence are incompletely understood. On the one hand, improved diagnostics and surveillance methods as well as increased awareness of whooping cough by health care professionals might contribute ^1^. On the other hand, molecular changes in the pathogenic *B. pertussis* lineages and waning immunity especially after receipt of acellular pertussis vaccines have been debated ^1^. These data highlight the need to improve the vaccine formulations and vaccination campaigns, but also to develop alternative means to treat whooping cough.

Pertussis toxin is a major virulence factor of *B. pertussis* ^6^, and a detoxified form of the toxin is included in all acellular pertussis vaccine formulations. Clinical isolates lacking pertussis toxin have turned out to be rare ^7^. When administrated systemically to experimental animals, e.g. to mice ^8^, pertussis toxin recapitulated the leukocytosis (increase in number of circulating white blood cells) seen in young children with whooping cough. Rats experimentally infected with *B. pertussis* developed prolonged paroxysmal coughing, but an isogenic pertussis toxin-deficient strain did not cause such pathology ^9, 10^. Seven-day-old neonatal mice infected with a pertussis toxin-deficient strain of *B. pertussis* survived a challenge, which caused 100% mortality with the parental strain ^11^. Therefore, targeting of pertussis toxin might prove beneficial in the treatment of whooping cough, especially in young children who still lack the vaccine-induced protection.

Pertussis toxin is composed of five non-covalently bound subunits (PtxS1-S5), which are arranged in an AB_5_-topology ^12, 13^. The B_5_-assembly is formed by the PtxS2-S5 subunits (PtxS2, PtxS3, 2 copies of PtxS4, PtxS5) ^12, 13^. Pertussis toxin is secreted from the bacteria via Sec-pathway and Ptl type IV secretion system ^14^. The B_5_-assembly mediates binding of the secreted AB_5_ holotoxin on the surface of various different cell types in a carbohydrate-dependent manner ^13^. Subsequent cell entry is followed by dissociation of the B_5_-assembly and the PtxS1-subunit ^15^, which belongs to the family of ADP-ribosyltransferases (ARTs) ^16^. PtxS1 ADP-ribosylates a single C-terminal cysteine residue in α-subunits of most heterotrimeric Gαi-superfamily members such as Gαi, Gαo, and Gαt ^17-19^. The C-terminus of the heterotrimeric G-protein α-subunits makes functionally important contacts with the plasma membrane-localized G-protein coupled receptors (GPCRs) ^20^. The bulky PtxS1-catalyzed ADP-ribose modification decouples the G-protein αi-subunit, and also the βγ-dimer, from the GPCRs and inhibits the agonist stimulation-induced signal propagation inside the cell.

In this study, we set out to identify small molecular weight compounds that inhibit the Gαi-specific ADP-ribosyltransferase activity of pertussis toxin, and thereby to potentially interfere with the pathological effects of pertussis toxin.

## RESULTS

### Purification of recombinant PtxS1 (rPtxS1) from *E. coli*

We did not succeed in purifying the full length N- or C-terminally HIS-tagged recombinant *B. pertussis* PtxS1 from *E. coli* due to weak solubility and proteolytic instability (data not shown). Therefore, we engineered a double deletion to *ptxA* resulting into expression of an N-terminally HIS-tagged version of *B. pertussis* Tohama I strain PtxS1 (UniProt_P04977), referred hereafter to as rPtxS1, which lacks the N-terminal secretion signal sequence as well as part of the C-terminus (Fig. 1, Fig. 2A). Structural data of the pertussis AB_5_ holotoxin demonstrate that the C-terminus of PtxS1 masks the active site, and indicate that di-sulfide bond reduction (Cys41-Cys201) and significant movement of the C-terminus would be required for catalysis^12, 13^. Our C-terminal truncation exposes the ADP-ribosyltransferase catalytic site of PtxS1 including the NAD^+^-binding pocket (Fig. 1)^12, 13^.

**Figure 1.**
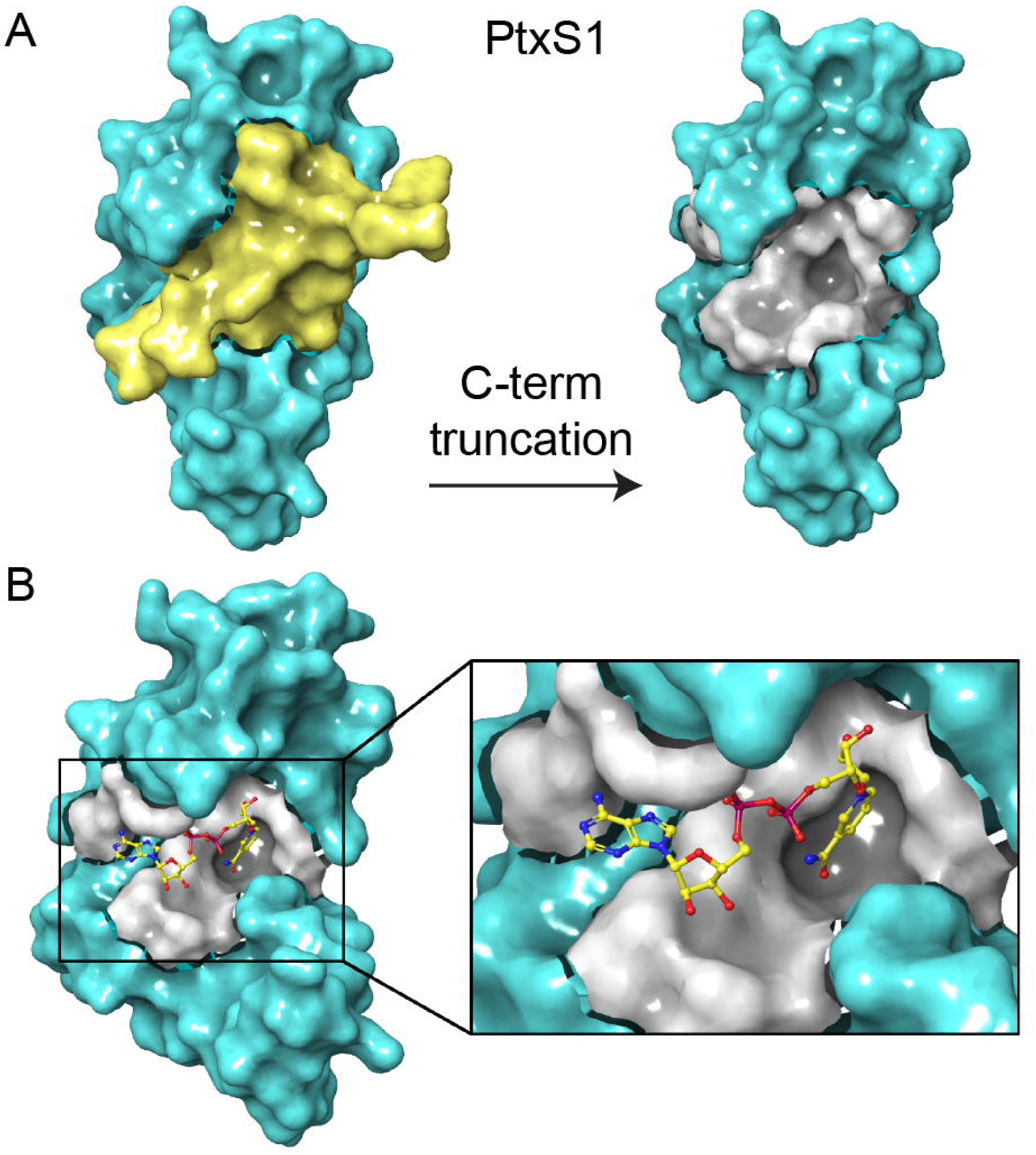
Ptx51 of pertussis toxin. **A)** Surface representation of the crystal structure of PtxS1 (PDB_1BCP). The truncated C-terminal fragment and the truncation-uncvoered ART active site are shown in yellow and grey, respectively. **B)** Possible binding mode of NAD^+^ to PtxS1 derived by superimposing the PtxS1 crystal structure (PDB_1BCP) to the crystal structure of active site mutant of pertussis-like toxin from *E. coli* with bound NAD^+^ (PDB_4Z9D). Color-coding of atoms in NAD^+^: yellow, carbon; blue, nitrogen; red, oxygen; magenta, phosphorus.

**Figure 2.**
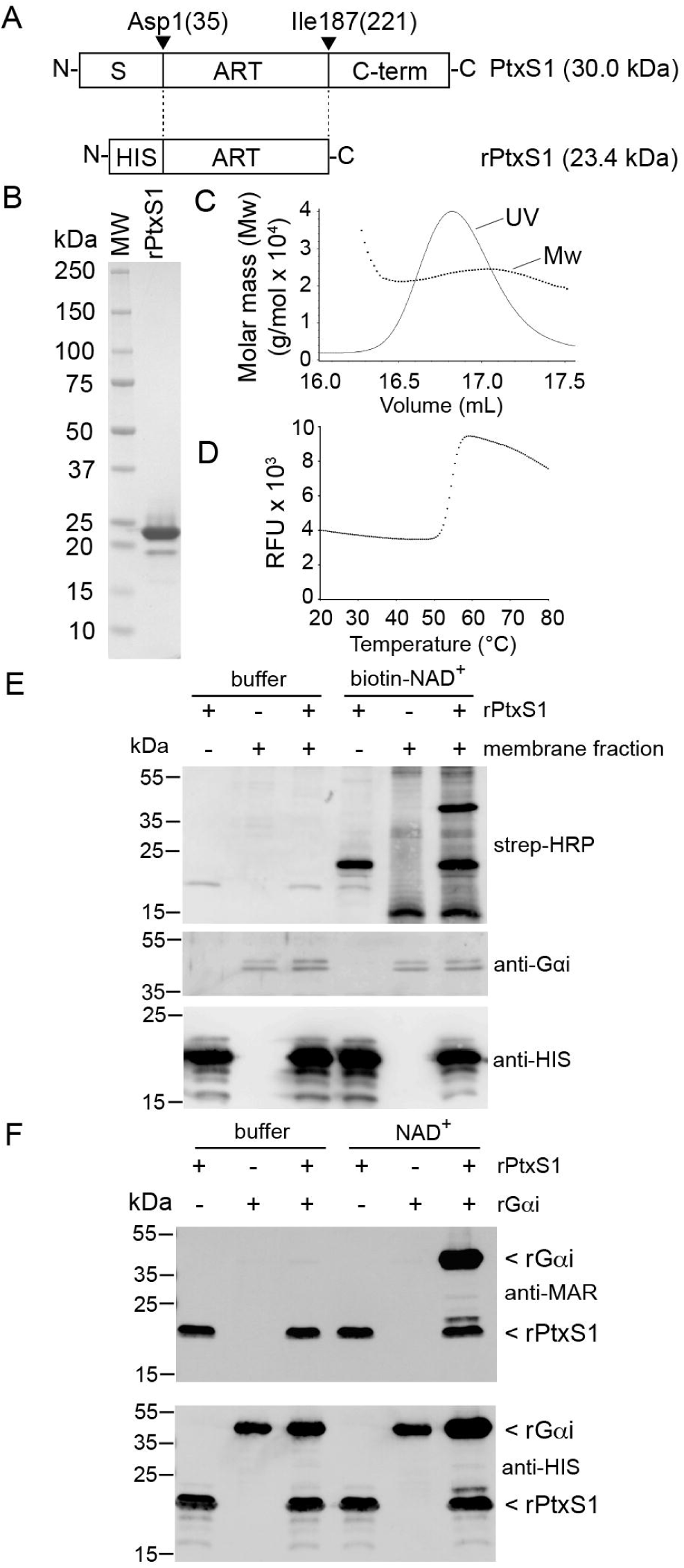
Catalytic activity of rPtxS1. **A)** Construct design. rPtxS1 lacks the N-terminal secretion signal (S) as well as part of the C-terminus (C-term). N-terminus of the mature PtxS1 starting with Asp35 is classically numbered as the first amino acid of PtxS1. **B)** PageBlue-stained SDS-PAGE gel of SEC-purified rPtxS1. **C)** Representative SEC-MALS result of SEC-purified rPtxS1. **D)** Representative DSF result of SEC-purified rPtxS1. **E)** *In vitro* ADP-ribosylation assay (enzyme-excess condition) for rPtxS1 with NAD^+^-biotin and HEK293T membrane fraction with endogenous level of Gαi. Protein-conjugated ADP-ribose-biotin is detected with streptavidin-HRP **F)** *In vitro* ADP-ribosylation assay (enzyme-excess condition) for rPtxS1 with NAO^+^ and recombinant N-terminally HIS-tagged Gαi. Protein-conjugated ADP-ribose is detected with a rabbit polyclonal antibody specific for mono-ADP-ribose. Same samples were analyzed in three (Fig. 2E) and two (Fig. 2F) parallel membranes, respectively.

rPtxS1, as purified with metal affinity and size exclusion chromatography (SEC), is proteolytically stable and migrates between the 25 and 20 kDa protein markers in SDS-PAGE (Fig. 2B), in accordance with a theoretical size of 23.4 kDa. In order to study the oligomerization state of rPtxS1 in solution, size exclusion chromatography coupled to multi-angle light scattering (SEC-MALS) was used. Purified rPtxS1 appeared as a single peak (Fig. 2C), with an estimate of 23.04 ± 0.2 kDa in a triplicate run indicating that rPtxS1 is monomeric in solution. rPtxS1 is well folded as evidenced by a sigmoidal curve with a melting temperature (Tm) of 54.67 ± 0.22 °C in a differential scanning fluorimetry (DSF)-assay (Fig. 2D).

### Analysis of the ART activity of rPtxS1

ART activity of rPtxS1 towards Gαi was first studied with a HEK293T cell membrane preparation having an endogenous level of Gαi in an NAD^+^-biotin Western blot-assay. rPtxS1 ADP-ribosylated one major protein from the complex membrane proteome (Fig. 2E). This protein migrated between the 55 and 35 kDa protein markers in SDS-PAGE (Fig. 2E), in accordance with a theoretical size of endogenous Gαi, e.g. 40.4 kDa for isoform 1 (UniProt_P63096). Next, ART activity of rPtxS1 towards Gαi was studied with recombinant N-terminally HIS-tagged Gαi (isoform 1, UniProt_P63096) purified from *E. coli*, hereafter referred to as rGαi, in an NAD^+^ Western blot-assay. rPtxS1 ADP-ribosylated rGαi (Fig. 2F), indicating that Gαi may efficiently serve as a substrate without the G-protein βγ-complex or other cellular constituents. SEC-analysis of rPtxS1-rGαi complex solution topology showed the presence of only single monomeric rPtxS1 and rGαi proteins (Fig. S1). rPtxS1-rGαi complex formation during catalysis therefore appears to be weak, i.e. the rGαi modification is based on a transient kiss-and-run interaction. Interestingly, rPtxS1 modified itself (Fig. 2E), and it already had become auto-ADP-ribosylated in the *E. coli* expression host (Fig. 2F). This activity was drastically diminished in a Q127D/E129D double mutant of rPtxS1(Fig. S2A), hereafter referred to as rPtxS1-mutant. Based on a DSF-assay, the Tm of the rPtxS1-mutant was 52.00 ± 0 °C as compared to Tm of the rPtxS1-wt (Tm of 54.67 ± 0.22 °C), indicating that the Q127D/E129D double mutation does not affect the folding of rPtxS1. The rPtxS1-mutant was also incapable of ADP-ribosylating rGαi (Fig. S2B). This double mutant of rPtxS1 was analyzed, because structurally identical mutations in a recently discovered pertussis-like toxin from *E. coli* caused catalytic inactivation, although NAD^+^ was still capable of binding to the protein (Fig. S2C) ^21^. Q127 and E129 of rPtxS1 are also conserved with several other bacterial ART-toxins where these residues position NAD^+^, in particular the nicotinamide end (Fig. S2C), and promote the transfer of ADP-ribose to a substrate amino acid residue ^16^. To assess the substrate amino acid specificity of the extensively truncated rPtxS1 (see Fig. 1 and Fig. 2A), we analyzed the C351A mutant of rGαi (UniProt_P63096), hereafter referred to as rGαi-mutant, in an NAD^+^-biotin Western blot-assay. rPtxS1 ADP-ribosylated rGαi, but it was incapable of ADP-ribosylating the rGαi-mutant (Fig. S3). Based on a DSF-assay, the Tm of the rGαi-mutant was 44.33 ± 0.23 °C as compared to Tm of the rGαi-wt (Tm of 43.83 ± 0.23 °C), indicating that the C351A mutation does not affect the folding of rGαi. Therefore, rPtxS1 is not only catalytically active towards rGαi, but it also retains the Gαi substrate amino acid specificity evidenced with the pertussis AB_5_ holotoxin ^17-19^.

### Multiwell-based assay set-up for screening of rPtxS1 inhibitors

We analyzed the suitability of a fluorometric method to screen for rPtxS1 inhibitors previously developed for poly-ADP-ribose (PAR) synthesizing enzymes ^22^ and later extended to mono-ADP-ribose (MAR) synthesizing enzymes ^23^. The end point assay is based on conversion of NAD^+^ to a stable fluorophore with emission maximum at 444 nm ^22^. The decrease in fluorescence in comparison to the non-enzyme control is a measure of NAD^+^-consuming enzymatic activity ^22^. Incubation of rPtxS1 with rGαi for 40 min resulted in a strong decrease of fluorescence (Fig. 3A). No fluorescence decrease was detected with the rPtxS1-mutant under identical conditions (Fig. 3A). Assays with different concentrations of rPtxS1 were stopped at various time points to measure NAD^+^ consumption in the presence of a constant amount of rGαi substrate. These data demonstrated that an increase in the rPtxS1 concentration increases NAD^+^ consumption that also progresses over time (Fig. 3B). rPtxS1 could also consume NAD^+^ in the absence of rGαi, although much slower than with the rGαi substrate (Fig. 3A). This NAD^+^ glycohydrolase activity, i.e. enzyme-catalyzed reaction between NAD^+^ and water to yield ADP-ribose and nicotinamide, was not detected with the rPtxS1-mutant (Fig. 3A). The effect of dimethyl sulfoxide (DMSO) on the NAD^+^-consuming activity of rPtxS1 was also studied because small molecules in chemical libraries are usually dissolved in DMSO typically resulting in assay concentrations of 0.1% – 1% DMSO. DMSO did not have a significant effect on the NAD^+^-consuming activity of rPtxS1 in the presence of rGαi substrate up to 0.2% DMSO, while 1% DMSO already significantly (*p* = 10^−4^) reduced activity (Fig. 3C). We chose to perform compound screening for rPtxS1 inhibitors in the presence of 0.1% DMSO. To test reproducibility of the fluorometric assay, maximum (NAD^+^ as incubated in plain buffer) and minimum (NAD^+^ as incubated in rPtxS1- and rGαi-containing buffer) signals were measured from five independent runs to test plate-to-plate and day-to-day variability. The average Z’ value for the assay was 0.68, indicating that the fluorometric assay is suitable for high throughput screening. The statistical parameters of the NAD^+^ quantitation assay are summarized in Table S1.

**Figure 3.**
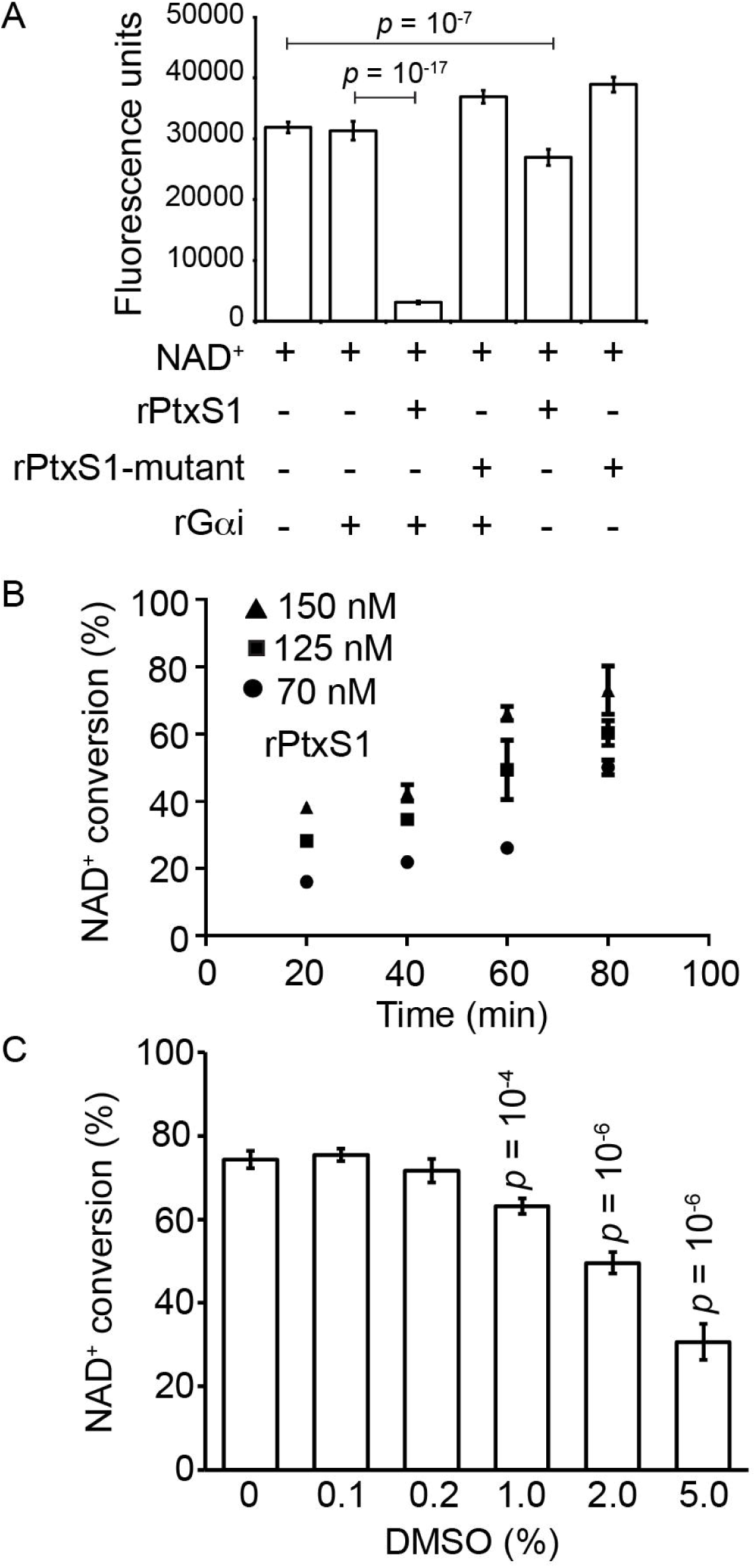
Multiwell fluorometric NAD^+^ quantitation assay. Decrease in fluorescence over time is a measure of NAD^+^-consuming enzymatic activity **A)** rPtxS1 and rPtxS1-mutant were incubated for 40 min in the presence or for 60 min in the absence of rGαi substrate. **B)** Time and concentration dependency of NAD^+^-consuming enzymatic activity of rPtxS1 in the presence of rGαi substrate. **C)** Effect of DMSO on NAD^+^-consuming enzymatic activity of rPtxS1 in the presence of rGαi substrate. Statistics by two-tailed Student’s t-test two sample equal variance.

### Screening for rPtxS1 inhibitors

We first analyzed a panel of selected compounds (n = 32) known to inhibit the ART-activity of human diphtheria toxin-like ADP-ribosyltransferases (ARTDs/PARPs). These compounds (Table S2), which also included several FDA-approved anti-cancer drugs such as Rucaparib, did not inhibit the NAD^+^-consumption activity of rPtxS1 in the presence of rGαi when analyzed at 80-fold molar excess to rPtxS1 (arbitrary inhibitory threshold = 50%). Next, we screened a diversity set compound library obtained from the National Cancer Institute (NCI) Developmental Therapeutics program repository (https://dtp.cancer.gov), which at the time of acquiring contained 1,695 compounds (diversity set III). The so-called diversity set contains selected compounds, e.g. to represent high scaffold diversity, from the over 100,000 NCI-repository compounds. A total of nine compound hits were identified, which inhibited the NAD^+^-consumption activity of rPtxS1 in the presence of rGαi more than 50% when analyzed at 80-fold molar excess to rPtxS1. These nine compounds were analyzed in an independent NAD^+^-biotin Western blot-assay, which measures the rPtxS1-catalyzed conjugation of ADP-ribose-biotin onto rGαi. Five out of the nine primary compound hits showed strong inhibition of rPtxS1-catalyzed rGαi ADP-ribosylation when analyzed at 200-fold molar excess to rPtxS1 (Fig. 4A, titration effect shown for NSC228155 and NSC29193 in Fig. 4B). NSC119875 (Cisplatin, DNA alkylating agent used in cancer treatments) and NSC44750 affected rGαi and/or rPtxS1 protein integrity in the NAD^+^-biotin Western-blot based assay (Fig. 4A), even if the assay was performed with a 10-fold lower compound concentration (data not shown). NSC119875 and NSC44750 were therefore excluded from further studies. To evaluate the potency of the remaining three compounds to inhibit rPtxS1, dose response studies were performed using the fluorometric NAD^+^-consumption assay in the presence of rGαi. NSC228155 and NSC29193 had IC_50_-values of 3.0 and 6.8 μM, respectively (Fig. 4C). NSC149286 was less potent than the other two compounds with an IC_50_-value of 20 μM (Fig. 4C), and was excluded from further analyses. Chemical structures of the hit compounds are shown in Figure 4C. NSC228155 and NSC29193 appeared to have mimicry to the adenine base or the nicotinamide end of NAD^+^ (Fig. 4D). We also utilized an alternative readout of the NAD^+^-biotin assay, i.e. capture of HIS-tagged proteins on 96-well nickel-coated plates followed by the detection of protein-conjugated ADP-ribose-biotin. As shown in Figure 4E, NSC228155 and NSC29193 showed strong inhibition of the rPtxS1-catalyzed conjugation of ADP-ribose-biotin onto rGαi. NSC228155 and NSC29193 also inhibited the auto-ADP-ribosylation activity of rPtxS1 (Fig. 4F), indicating that these two compounds interact directly with rPtxS1. The two hit compounds were not identified as pan-assay interference compounds or aggregators (zinc15.docking.org). In summary, compound screening resulted into identification of NSC228155 and NSC29193, which inhibited the ART activity of rPtxS1 in low micromolar concentrations.

**Figure 4.**
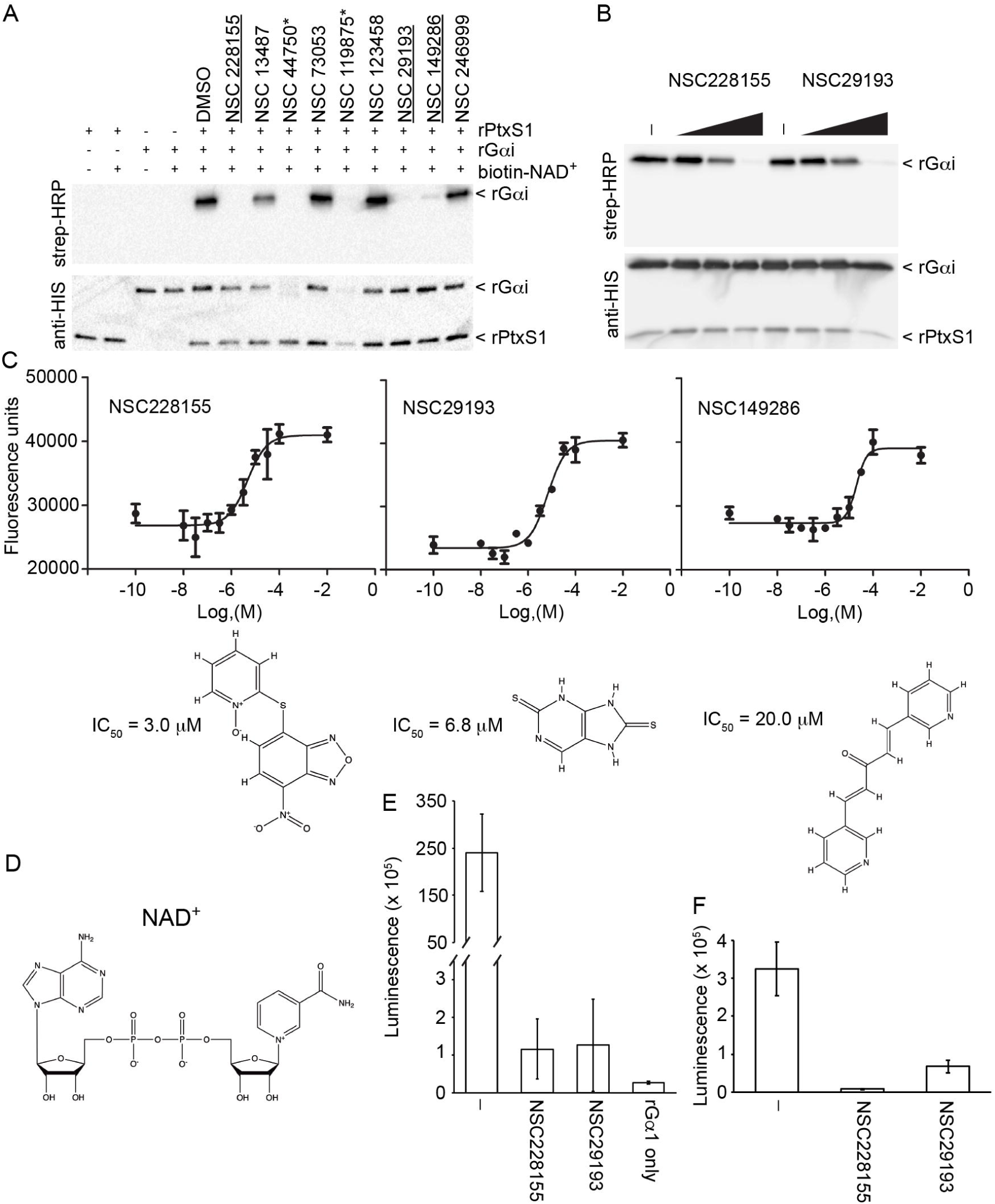
Evaluation of primary compound hits from rPtxSI inhibitor screen. **A)** *In vitro* ADP-ribosylation assay (substrate-excess condition) for rPtxSI with NAD^+^-biotin and rG αi. Protein-conjugated biotin-ADP-ribose was detected with streptavidin-HRP. The reactions contained 200-fold molar excess of inhibitors over rPtxSI. Compounds marked with an asterisk affect rPtxSI and/or rG αi protein integrity, and were therefore excluded from further studies. Same samples were analyzed in two parallel membranes. **B)** *In vitro* ADP-ribosylation assay (substrate-excess condition) for rPtxSI with NAD^+^-biotin and rG αi (see also Fig. 4A). The reactions contained 2-, 20-, or 200-fold molar excess of inhibitors over rPtxSI. Same samples were analyzed in two parallel membranes **C)** IC_50_-curves and chemical structures of the hit compounds selected based on Fig. 4A data. **D)** Chemical structure of NAD*. **E-F)** In vitro ADP-ribosylation assay in a 96-well nickel-coated plate format. Protein-conjugated biotin-ADP-ribose (E: rPtxSI + rG αi or rG αi only reaction conditions; F; rPtxSI only reaction condition) was detected with streptavidin-HRP. The reactions contained 400-fold molar excess of inhibitors over rPtxSI.

### Putative binding poses of NSC29193 and NSC228155 to PtxS1

In order to evaluate possible binding modes of the compounds to PtxS1, we performed docking and molecular dynamic simulation (MDS). The top ranked poses of NSC228155 shared features, including the placement of the aromatic pyridine ring of NSC228155 in an enclosed region where the nicotinamide ring of NAD^+^ would bind to PtxS1, with the double benzoxadiazole ring positioned near the center of the NAD^+^-binding pocket. The benzoxadiazole ring is joined to the pyridine ring via a central sulfur atom, and the rotatable bonds permit the adoption of varying conformations. In pose 1 (Fig. 5A), with the lowest free energy of binding (ΔG_bind_ of −51.2 kcal/mol; docking score of −4.08 kcal/mol), the pyridine ring oxygen of the ligand forms a hydrogen bond with the main-chain nitrogen atom of Tyr10, anchoring the ligand to an aromatic area of the binding pocket, and with π-π stacking interactions observed for the pyridine ring with nearby Tyr59 and for the benzoxadiazole ring with Tyr63. In pose 2 (Fig. 5B), having the best docking score (−4.20 kcal/mol; ΔG_bind_ of −43.3 kcal/mol), the pyridine ring is rotated 90° relative to pose 1, losing the backbone hydrogen bond with Tyr10, but making π-π interactions with both Tyr63 and Tyr10, and ionic interactions with Arg9 and Arg67. In MDS, pose 1 showed stability and retained its position with respect to the original docked position, only differing on average by an root-mean-square deviation (RMSD) of 0.9 Å (Fig. S4A), and the RMSD for the ligand orientations differ on average by 2.3 Å when the coordinates of PtxS1 are superposed over the backbone atoms. In contrast, pose 2 was unstable at the initial binding position and occupied different locations of the binding pocket with an average 22 Å RMSD for the ligand based on superposing the PtxS1 coordinates from the simulation (Fig. S4B).

**Figure 5.**
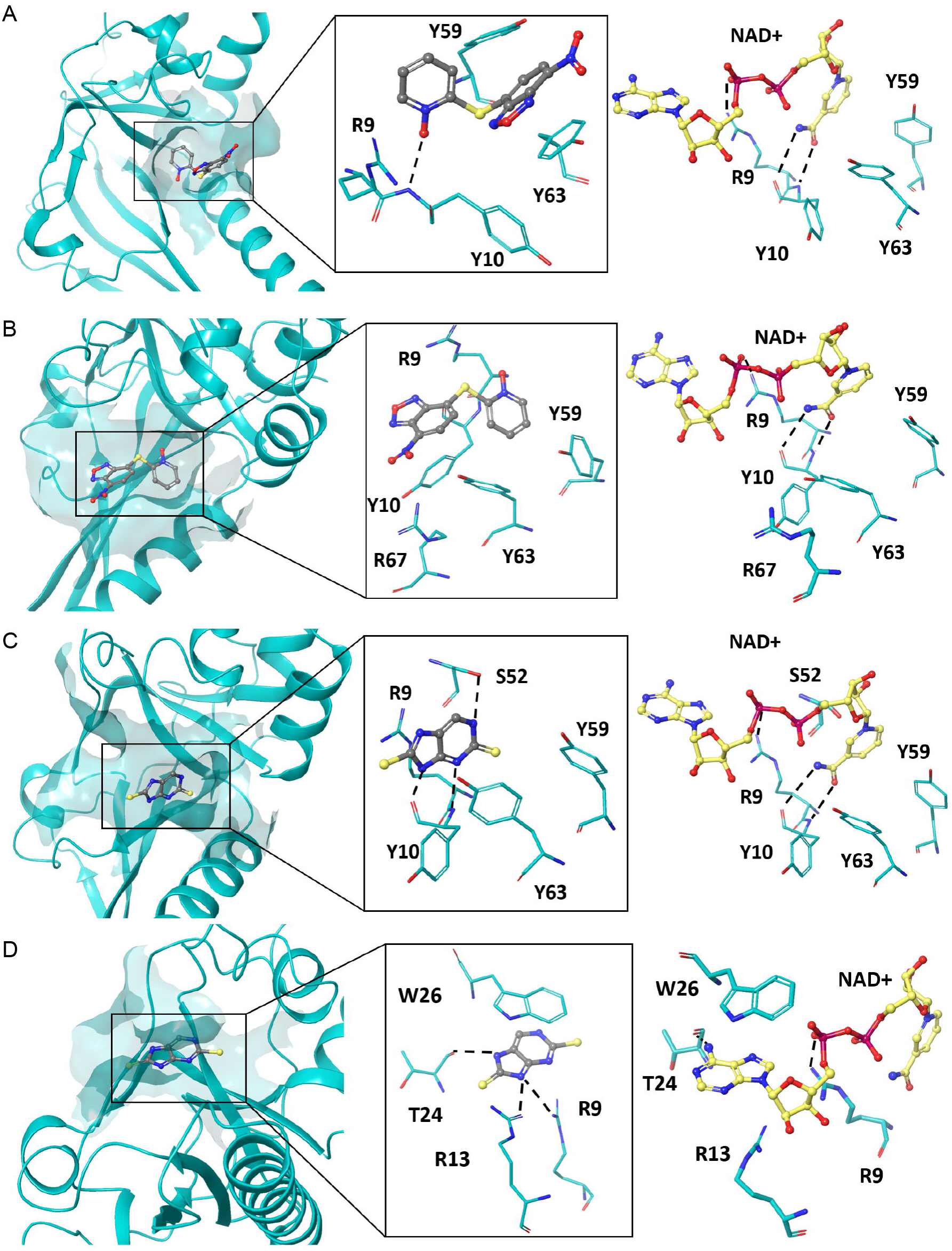
Prediction of binding poses of NSC228155 and NSC29193 to Ptx51. Selected binding poses 1 **A)** and 2 **B)** of NSC228155 and binding poses 1 **C)** and 2 **D)** of NSC29193 to PtxS1 (POB_1BCP, chain A), and residues involved in ligand interactions. Binding mode of NAO^+^ to PtxS1 in the corresponding area of the NAO^+^-binding pocket is shown in each panel on the right (see also Fig. 1). Color-coding of atoms in NAO^+^: yellow, carbon; blue, nitrogen; red, oxygen; magenta, phosphorus. Hydrogen bond interactions are shown in dotted lines.

In contrast to NSC228155, NSC29193 is rigid and almost exclusively resulted in two docked poses. Pose 1 of NSC29193 is positioned near the binding site for the nicotinamide ring of NAD^+^in the PtxS1, hydrogen bonding with main-chain nitrogen and oxygen atoms of Tyr10 (Fig. 5C). Additionally, NSC29193 forms π-cation interactions with Arg9, π-π stacking with Tyr63 and hydrogen bonding with the side-chain hydroxyl group of Ser52. NSC29193, pose 2, is bound near the location where the adenine ring of NAD^+^ would be bound in the pertussis toxin, stacking against the aromatic ring of Trp26 and hydrogen bonding with the main-chain oxygen atom of Thr24 (Fig. 5D) as well as forming ionic interactions with Arg9 and Arg13. During MDS, NSC29193 pose 1 was quite stable, with an average RMSD of 0.1 and 1.0 Å with respect to the ligand and receptor backbone atoms (Fig. S4C), and interactions between the ligand and main-chain atoms of Tyr10 were conserved for over 90% of the simulation time. The π-π interactions with Tyr63 and π-cation interactions with Arg9 were also retained for the majority of the simulation time. In contrast, the binding mode of pose 2 was disrupted at around 20 ns of the simulation (Fig. S4D), mainly due to the reorientation of the indole ring of Trp26, which appears to be one of the key interaction partners of NSC29193. Consequently, in pose 2 the ligand moves from the initial binding position and binds to several areas of the binding pocket. This is reflected by the average 12 Å RMSD of the ligand with respect to the superposed coordinates of the toxin from the simulation (Fig. S4D). Taken together, docking and MDS resulted in identification of plausible binding poses of NSC228155 and in particular for NSC29193, most likely owing to the rigidity of the latter ligand, at the NAD^+^-binding pocket of PtxS1.

### NSC228155 and NSC29193 as inhibitors of pertussis toxin in living human cells

First, to find the optimal toxin dosage, we titrated the pertussis AB_5_ holotoxin and detected the ADP-ribosylation of endogenous Gαi by Western blot-assay using a polyclonal MAR-recognizing antibody. Incubation of HEK293T cells for 2 h with the holotoxin resulted, in a concentration dependent-manner, in mono-ADP-ribosylation of a single protein migrating between the 35 and 55 kDa protein markers in SDS-PAGE (Fig. 6A), in accordance with a theoretical size of endogenous Gαi, e.g. 40.4 kDa for isoform 1 (UniProt_P63096). Next, we pre-incubated the HEK293T cells for 30 min with the compounds NSC228155 or NSC29193 prior to addition of the holotoxin and subsequent co-incubation. We did not detect inhibitory action for the compound NSC29193 despite multiple analyzed conditions (data not shown), e.g. i) 30 min pre-incubation with inhibitor (0.1, 1, 10 and 50 µM) + 2 h with holotoxin (10 ng/mL) or ii) 1, 2, 3, 4 and 5 h pre-incubation with 50 µM inhibitor + 2 h with holotoxin (10 ng/mL). However, compound NSC228155 inhibited the Gαi-specific ADP-ribosylation activity of holotoxin in a concentration-dependent manner (Fig. 6B). Some inhibitory action was evidenced with 0.1 and 1 µM of NSC228155, but 5 µM of NSC228155 resulted in a near complete inhibition of mono-ADP-ribosylation of Gαi. Based on pixel intensity analysis of the anti-MAR signal, the IC_50_ of NSC228155 inhibitory effect was 2.35 µM (Fig. 6C).

**Figure 6.**
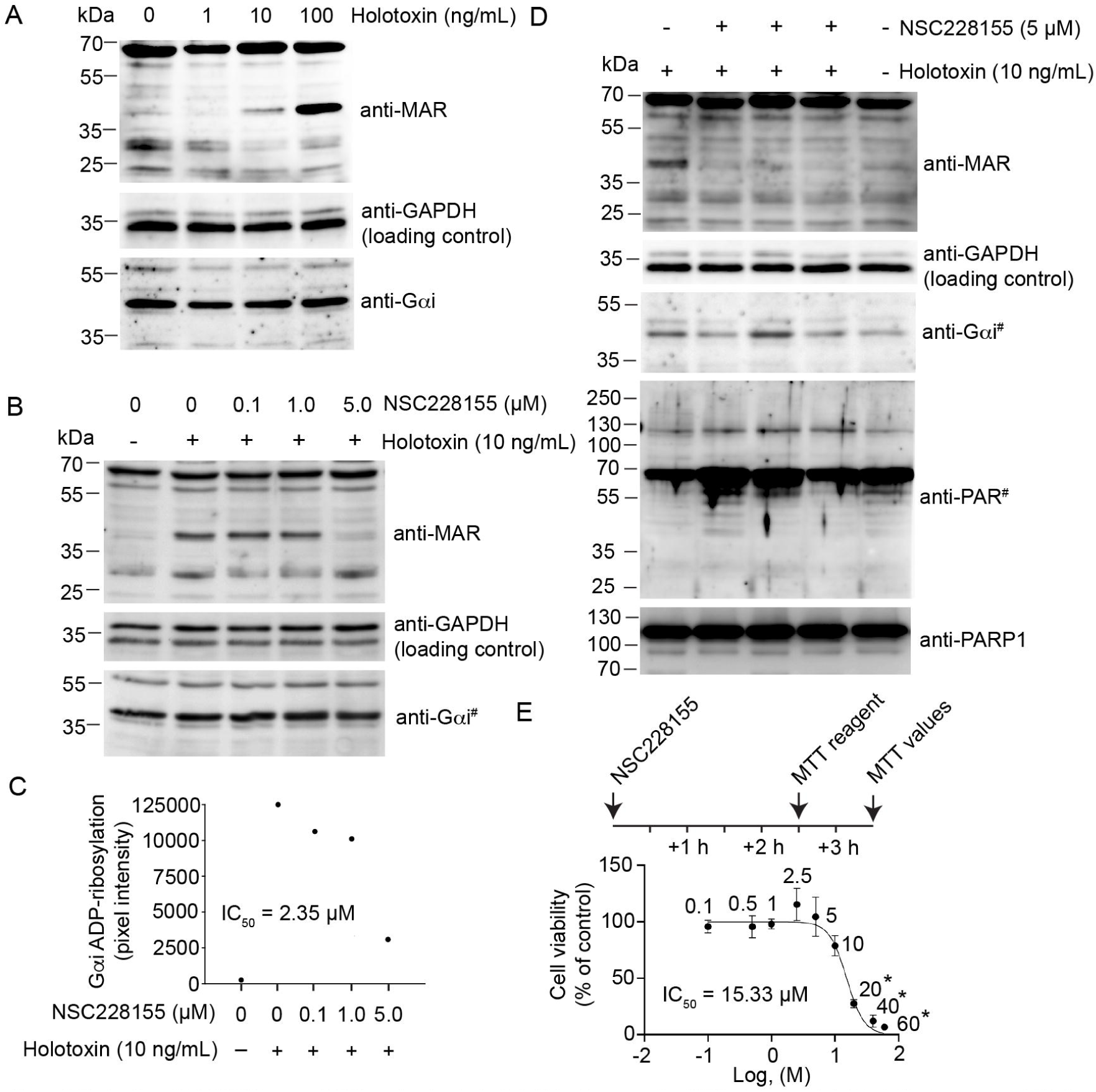
Evaluation of NSC228155 as an inhibitor of pertussis toxin in living human cells. **A)** Effect of pertussis holotoxin titration on mono-ADP-ribo-sylation of Gαi in living HEK293T cells (2 h incubation). Protein-conjugated ADP-ribose is detected with a rabbit polyclonal antibody specific for mono-ADP-ribose (MAR). **B)** Effect of NSC228155 titration on pertussis toxin-catalyzed mono-ADP-ribosylation of Gαi in living HEK293T cells. Inhibitors were added 30 min before starting the 2 h holotoxin incubation. “Western blot of a parallel membrane with the same Fig. 6B samples, otherwise Fig. 6A and Fig. 6B blots probed, stripped and re-probed in the order of 1) anti-MAR, 2) anti-GAPDH and 3) anti-Gαi. **C)** Quantitation of the inhibitory effect of NSC228155 on pertussis holotoxin-catalyzed mono-ADP-ribosylation of Gαi. Pixel intensities of the anti-MAR signal (Fig. 6B) were normalized based on the anti-GAPDH loading control. **D)** Effect of 5 pM NSC228155 (triplicate treatment) on pertussis holotoxin-catalyzed mono-ADP-ribosylation of Gαi and cell viability as assessed by the DNA damage-associated auto-poly-ADP-ribosylation of PARP1 and apoptosis-associated PARP1 proteolytic processing. #Western blot of a parallel membrane with the same Fig. 6D samples, otherwise Fig. 6D blots probed, stripped and re-probed in the order of 1) anti-MAR, 2) anti-GAPDH and 3) anti-PARP1. **E)** Effect of NSC228155 titration on metabolic activity of HEK293T cells as analyzed by the MTT-assay 3.5 h after compound addition. Statistics by two-tailed Student’s t-test two sample equal variance (*p-value <0.05).

Having established NSC228155 as a potent inhibitor of pertussis toxin in living HEK293T cells, we turned our attention to the maximum tolerated dose. NSC228155, at the effective 5 µM inhibitory concentration, did not significantly increase the DNA-damage induced PARP1 (also called ARTD1) poly-ADP-ribosylation or cell death-associated PARP1/ARTD1 proteolytic processing from the approximately 120 kDa full length form into the approximately 90 kDa form (Fig. 6D). However, visual alterations, including partial cell detachment, was detected in HEK293T cell monolayers upon titration of NSC228155 up to 50 µM in the 2.5 h Gαi ADP-ribosylation assay (data not shown). Therefore, we utilized the MTT-assay, i.e. reduction of tetrazolium salt into formazan by metabolically active cells, to study the potential cytotoxicity of NSC228155. The metabolic activity of HEK293T cells, as analyzed by the MTT-assay 3.5 h after addition of NSC228155, was significantly affected with 20 µM or higher concentrations (Fig. 6E). IC_50_ of NSC228155 cytotoxicity was 15.33 µM (Fig. 6E). In conclusion, NSC228155 is a potent inhibitor of pertussis toxin in living HEK293T cells, but at high concentrations it induces cytotoxicity. The mechanism(s) of NSC228155 cytotoxicity needs to be addressed in the future hit development of NCS228155.

## DISCUSSION

We report the identification of the first small molecular weight compounds inhibiting the ADP-ribosyltransferase activity of pertussis toxin *in vitro* and in living cells. ADP-ribosyltransferase activity of pertussis toxin is important for the pathological effects of pertussis toxin. Seven-day-old neonatal mice infected with a *B. pertussis* strain expressing a mutant of PtxS1 lacking the ART-activity fully survived a challenge, which caused 100% mortality with the parental strain of *B. pertussis* ^11^. Purified AB_5_ holotoxin containing the same PtxS1 mutant was incapable of inducing leukocytosis in mice in doses that were 100-fold more than the median lethal dose of wild-type AB_5_ holotoxin ^24^. Therefore, targeting of the ADP-ribosyltransferase activity of pertussis toxin might prove beneficial in the treatment of whooping cough.

Compound library screening was made possible due to our ability to purify mg-levels of a truncated but highly active recombinant *B. pertussis* PtxS1 (rPtxS1) from *E. coli*. rPtxS1 retained the high amino acid specificity of pertussis AB_5_ holotoxin toward the single C-terminal cysteine in Gαi ^17-19^. Used truncation positions in rPtxS1 are functionally justified. PtxS1 contains an N-terminal signal sequence ending at Ala34 ^25, 26^ that gets cleaved upon the Sec-mediated secretion of PtxS1 from the cytoplasm into the periplasm where the AB_5_ holotoxin is assembled prior to Ptl type IV secretion system-mediated export. The truncated part of the PtxS1 C-terminus masks the NAD^+^-binding pocket in the AB_5_ holotoxin involving a stabilizing intramolecular di-sulfide bond between Cys41 and Cys201 ^12, 13^. Our truncation approach bypasses the need for disulfide bond reduction ^27^ and extensive conformational movement of the C-terminus ^12, 13^ to activate PtxS1, which inside the host cell has also been proposed to involve proteolysis ^28^.

We made an initial attempt to identify PtxS1 inhibitory compounds from molecules (n = 32, see Table S2) known to inhibit the ART-activity of human diphtheria toxin-like ADP-ribosyltransferases (ARTDs/PARPs). We envisioned that some of these compounds, which target the ARTD/PARP NAD^+^-binding pockets, could provide a fast track to antibiotic development via repurposing drug strategy, due to the current FDA-approvals in cancer therapy. However, none of these compounds significantly inhibited the NAD^+^-consumption activity of rPtxS1 in the presence of rGαi. Therefore, we decided to screen a diversity set compound library (n = 1,695) obtained from the National Cancer Institute (NCI) Developmental Therapeutics program repository (https://dtp.cancer.gov). Two potent inhibitory compounds NSC29193 and NSC228155 with low micromolar IC_50_-values were identified in the *in vitro* NAD^+^ consumption assay. These compounds were also potent in an independent *in vitro* assay where we monitored the amount of rGαi-conjugated ADP-ribose-biotin upon rPtxS1 catalysis by streptavidin-HRP Western blotting (see Fig. 4B). NSC228155 and NSC29193 also inhibited the auto-ADP-ribosylation activity of rPtxS1 (see Fig. 4F), indicating that these two compounds interact directly with rPtxS1. Our docking and molecular dynamic simulations resulted in plausible binding poses for both ligands NSC228155 and NSC29193 (see Fig. 5 and S4). More consistent, energetically favorable binding poses were observed for NSC29193. This appears to be in accordance with the compound structures (see Fig. 4C). NSC29193 (purine-2,8-dithiol) is a rather rigid purine analogue that mimics the structure of the adenine base of NAD^+^ (see Fig. 4D). NSC228155, on the other hand, contains two major molecular structures, connected by a rotatable linker, one ring compound mimicking the adenine base of NAD^+^ and the other ring compound mimicking the nicotinamide of NAD^+^. It is noteworthy that 7 other compounds were also positive in the *in vitro* NAD^+^ consumption assay, some apparently via deleterious effects for protein stability as evidenced by the independent *in vitro* ADP-ribose conjugation assay (see Fig. 4A), i.e. compounds NSC44750 and NSC119875 (cisplatin). The data highlights the importance of validating the primary screening hits in alternative and independent *in vitro* assays. In summary, compound screening resulted into the identification of two compounds, NSC29193 and NSC228155, which inhibited the rPtxS1-catalyzed ADP-ribosylation of rGαi *in vitro* with low micromolar IC_50_-values.

NSC228155, but not NSC29193, was a potent inhibitor of pertussis holotoxin-mediated ADP-ribosylation of Gαi in living HEK293T cells. We analyzed multiple different experimental set-ups with NSC29193, including concentrations up to 50 µM that did not cause visible alterations or cell detachment of HEK293T cell monolayers. In respect of previous drug screening approaches there appears to be no published information on NSC29193. Accordingly, the cell permeability of NSC29193 is not known, but might be weak and thus explain our negative inhibitory data. In sharp contrast, we detected a near complete inhibition of the Gαi ADP-ribosylation with 5 µM NSC228155 (see Fig. 6). In part, this could relate to the fact that NSC228155 easily permeates cells. It has been shown in MDA-MB-468 breast cancer cells that 5 min incubation with 100 µM NSC228155 caused rapid movement of this inherently fluorescent molecule across cell membranes and dispersal to both the cytoplasm and nucleus ^29^. However, we witnessed significant toxicity of NSC228155 for HEK293T cells with 20 µM or higher concentrations after a 3.5-h incubation (cytotoxicity IC_50_ – 15.33 µM see Fig. 6E). This cytotoxicity might be caused by the proposed NSC228155-mediated production of reactive oxygen species (ROS) inside the cell ^29^. In this respect, it is noteworthy that, upon incubation of cells with 5 µM of NSC228155 for 2.5 h, we did not detect auto-ADP-ribosylation of PARP1/ARTD1 (see Fig. 6D), which is induced in cells upon ROS-induced DNA damage ^30^. Moreover, we did not detect caspase-mediated proteolytic PARP1/ARTD1 cleavage (see Fig. 6D), which is a robust readout for the onset of programmed cell death. However, future hit development needs to address the apparent cell toxicity effects of NCS228155 at high concentrations. Taken together, NSC228155 is a potent cell permeable hit compound to inhibit pertussis holotoxin-mediated ADP-ribosylation of Gαi in living cells in low micromolar concentrations.

## CONCLUSION

Resurgence of whooping cough has been witnessed even in highly vaccinated populations ^1-3^, and currently there are no specific therapeutics to treat whooping cough. Macrolide antibiotics, when administrated at a very early stage, show therapeutic effects in certain patient subgroups such as in infants <3 months of age ^31^. However, macrolide resistant *B. pertussis* strains have been reported ^4, 5^. Pertussis toxin is a major virulence factor of *B. pertussis* ^6^. Targeting of pertussis toxin might prove beneficial in the treatment of whooping cough, especially in young children who still lack the vaccine-induced protection against whooping cough. In this respect, several drug modes of action could be envisioned, including to inhibit – i) secretion from the bacterium, ii) host cell recognition, iii) endocytosis, iv) intracellular trafficking, and v) ADP-ribosylation activity inside the host cell. Humanized monoclonal antibodies have been developed that block pertussis toxin cell surface receptor interaction or the subsequent internalization and retrograde trafficking ^32^. These humanized antibodies prevented the characteristic signs of whooping cough in mouse and baboon models ^33^. Our current study shows that small molecular weight compounds inhibiting the ADP-ribosyltransferase activity of pertussis toxin might also have therapeutic potential. Similar findings have been published on some other ADP-ribosylating bacterial toxins, ExoS of *Pseudomonas aeruginosa* and cholix-toxin of *Vibrio cholerae* ^34, 35^. We conclude that NSC228155 and NSC29193 are useful templates for future hit development to specifically inhibit pertussis toxin ADP-ribosyltransferase activity in whooping cough.

## EXPERIMENTAL SECTION

### Expression plasmids

***i) rPtxS1-wt*** Synthetic DNA fragment (Eurofins Genomics) encoding for amino acids D35-I221 of *Bordetella pertussis* strain Tohama I (UniProt_P04977) was cloned with NdeI and BamHI into pET15b (Novagen) allowing expression of an N-terminally HIS-tagged rPtxS1 (MGSSHHHHHHSSGLVPRGSHM-D35-PtxS1-I221). The D35-I221 truncation positions of PtxS1 are based on ^36^. Of note, Asp35 is classically numbered as the first amino acid of PtxS1. ***ii) rPtxS1-Q127D/E129D*** pET15b-rPtxS1 plasmid was linearized with PCR using 5’-phosphorylated oligonucleotide primers (Eurofins Genomics) prAPV-351 (<u>GAT</u>AGCGA<u>T</u>TATCTGGCACACCGGCGCATTCCG, mutagenic nucleotides underlined) and prAPV-352 (GTAGGTGGCCAGCGCGCCGGCGAGGATACG). The PCR product was gel-isolated, re-ligated and transformed to acquire the mutant plasmid. ***iii) rG***α***i-wt*** Synthetic DNA fragment (GenScript) encoding for *E. coli* codon-optimized full-length human Gαi (isoform 1, UniProt_P63096-1) was cloned into pNIC-Bsa4 (Structural Genomics Consortium) using the ligation independent cloning method allowing expression of an N-terminally HIS-tagged rGαi (MHHHHHHSSGVDLGTENLYFQS-Gαi). ***iii) rG***α***i-Cys351Ala*** rGαi-encoding plasmid was used as a template in PCR to amplify Cys351Ala-mutant-encoding mutant allele using oligonucleotide primers (Eurofins Genomics) prAPV-418 (tacttccaatccATGGGTTGCACCCTGAGCGCGGAA, LIC-cloning overhangs in lowercase) and prAPV-419 (tatccacctttactgTCAGAACAGGCC<u>CGC</u>ATCCTTCAGGTTGTTCTTG, mutagenic nucleotides underlined, LIC-cloning overhangs in lowercase). The PCR product was cloned into pNIC-Bsa4 (Structural Genomics Consortium) using the ligation independent cloning method allowing expression of an N-terminally HIS-tagged rGαi-Cys351Ala mutant similar to rGαi-wt. All expression plasmids were verified by sequencing.

### Protein expression and purification

Expression plasmids were transformed into BL21(DE3) (Novagen) and selected overnight at 37°C on Luria-Bertani (LB) agar with appropriate antibiotics. Next morning, the bacterial lawn from the LB-plates was transferred into Terrific broth autoinduction medium (Formedium, AIMTB0205) supplemented with 0.8% (w/v) of glycerol with appropriate antibiotics. Cultures were grown at 37°C with 250 rpm until optical density at 600 nm reached 1 (typically 3-5 h), and temperature was reduced to 18°C. Bacteria were collected after 24 h by centrifugation and were either frozen to −80°C as pellets, in lysis buffer or directly used for purification. Pefabloc protease inhibitor (Roche, 11585916001) was added to 0.1 mM in the thawed biomass in lysis buffer [100 mM Hepes (pH 7.5), 500 mM NaCl, 10% (w/v) glycerol, 0.5 mM Tris(2-carboxyethyl)phosphine hydrochloride (TCEP), 10 mM imidazole]. Samples were sonicated and clarified by centrifugation. Supernatant was loaded to 5 mL HisTrap HP column (GE Healthcare). With rPtxS1 proteins, column was washed with 10 column volumes of wash buffer I and II. Wash buffer I and II have identical compositions to lysis buffer with the exception of imidazole concentration of 25 and 50 mM, respectively. rPtxS1 proteins were eluted with elution buffer [100mM Hepes (pH 7.5), 500 mM NaCl, 10% (w/v) glycerol, 0.5 mM TCEP, 500 mM imidazole], concentrated using a 10 kDa cut-off concentrator (Thermo Scientific) and subjected to size exclusion chromatography on Superdex75 16/600 Hiload Superdex column (GE Healthcare) using SEC buffer [100 mM Hepes (pH 7.5), 500 mM NaCl, 10% (w/v) glycerol, 0.5 mM TCEP]. With Gαi proteins, HisTrap HP column was washed with 15 column volumes of wash buffer [20mM Hepes (pH 7.5), 500 mM NaCl, 10% (w/v) glycerol, 0.5 mM TCEP, 25 mM imidazole] and eluted with elution buffer [20 mM Hepes (pH 7.5), 500 mM NaCl, 10% (w/v) glycerol, 0.5 mM TCEP, 500 mM imidazole]. Fractions were concentrated using a 10 kDa cut-off concentrator (Thermo Scientific) and further purified by size exclusion chromatography on Superdex75 16/600 Hiload Superdex column (GE Healthcare) with SEC buffer [30 mM Hepes (pH 7.5), 350 mM NaCl, 1 mM MgCl_2_, 0.5 mM TCEP]. Protein fractions were pooled, concentrated using a 10 kDa cut-off concentrator (Thermo Scientific), flash frozen and stored in −80°C.

### Multi-angle light scattering

All experiments were conducted with SEC-MALS buffer [100 mM Hepes (pH 7.5), 500 mM NaCl, 10% (w/v) glycerol, 0.5 mM TCEP] with a flow rate of 0.150 mL per minute. Buffer was filtered with 0.1 μm filter to remove small particles. Typically 100 µg of protein samples were injected into Superdex 200 10/300 increase column (GE Healthcare) by a Shimadzu autosampler coupled HPLC machine. Light scattering data were recorded using a multiangle light scattering detector (miniDAWN TREOS, Wyatt technology). Data were analyzed using Astra software (Wyatt technology). For complex formation studies, 100 µg of rPtxS1 and 100 µg of rGαi was incubated on ice for 2 h and then injected into the column.

### Differential scanning fluorimetry (DSF)

rPtxS1 at a concentration of 0.25 mg/mL was used in PBS. Protein was incubated with 5 × SYPRO orange (Thermo Scientific) for 10 minutes. The samples were heated from 20-90°C with 1°C increments (1 min/1°C). The experiment was run on 2500 Real-Time PCR systems (Applied Biosystems). The resulting data was analyzed with Boltzmann sigmoidal equation using GraphPad (GraphPad software, Inc.). Thermal stability of rGαi, rGαi-mutant and rPtxS1 alone as well as in the presence of NSC228155 or NSC29193, was analyzed by DSF using a CFX96 Real-Time PCR detection system (Bio-Rad). rGαi, rGαi-mutant and rPtxS1 at a concentration of 0.2 mg/mL was used in 20 mM Hepes (pH 7.5), 500 mM NaCl, 10% (w/v) glycerol, 0.5 mM TCEP. The inhibitor concentrations used in the assay ranged from 50 µM to 1 mM. Samples were incubated with 5 × SYPRO Orange (Thermo Scientific) for 5 min. The samples were heated from 20-90°C with 0.5°C increments (1 min/1°C). Tm-values were determined by using the CFX96 Real-Time PCR detection system (Bio-Rad) software.

### *In vitro* ADP-ribosylation assays

***i) enzyme-excess condition with Western blot read-out*** Reactions (typically in 120 µL) contained 10 μM rPtxS1 proteins, 10 μM biotinylated NAD^+^ (Trevigen, 4670-500-01) or 10 μM NAD^+^ (Sigma, N3014) and either 4 μM rGαi proteins or membrane fraction of HEK293T cells (30 µg of total protein) as the substrate in 100 mM Hepes (pH 7.5), 500 mM NaCl and 10% (w/v) glycerol. The membrane fraction was prepared essentially as described in ^37^. 80% confluent 10 cm cell culture plate of HEK293T cells was placed on ice and washed twice with PBS. Cells were collected by scraping into 1 mL of hypotonic lysis buffer [20 mM Hepes (pH 7.5), 2.5 mM MgCl_2_, 1 mM DTT supplemented with Pierce Protease and Phosphatase Inhibitor Mini Tablets (40 μL/mL of stock solution – one tablet / 2 mL H_2_O, Thermo Scientific, 88668) and 25 U/mL of benzonase (Merck-Millipore, 70664-3)]. The cells were incubated at 4°C for 1 h in rotation to allow them to swell and partially lyse. The partial lysates were drawn 20 times through 27-gauge needles and centrifuged with low speed (600 × g, 4°C, 10 min) to pellet the nuclei and insoluble cell debris. The low-speed supernatant was subjected to high-speed centrifugation (16100 × g, 4°C, 30 min) to pellet the membranes. The membranes were resolubilized into 50 μL of 50 mM Hepes (pH 7.5), 200 mM NaCl, 1 mM EDTA and 10 mM DTT, 0.3% (w/v) SDS and 2% Triton X-100 supplemented with protease and phosphatase inhibitors, in concentration as described above. Protein concentration was measured with Bradford assay. The ADP-ribosylation reactions were carried out at room temperature for 3 h with shaking at 300 rpm. Reactions were stopped by addition of Laemmli loading dye to 1 × and heating for 10 minutes at 95°C. The samples were run on SDS-PAGE and transferred to nitrocellulose membranes, followed by blocking with 1% (w/v) casein blocking buffer (Bio-Rad, 161-0782). Membranes were incubated with streptavidin conjugated to horse radish peroxidase (1:5000) (GE Healthcare, RPN1231VS) in 1% (w/v) casein blocking buffer (Bio-Rad, 161-0782) for 3 h at 4°C in rotation and washed thrice with Tris-buffered saline [10 mM Tris-HCl (pH 7.5), 150 mM NaCl] containing 0.05% Tween 20 (TBST) for ten minutes each time. Alternatively, after blocking with 4% (w/v) bovine serum albumin (BSA) in TBST, membranes were probed in TBST containing 2% (w/v) BSA (24 - 48 h at 4°C in rotation) for HIS-tagged rPtxS1 and rGαi proteins with mouse monoclonal anti-HIS (1:1000) (R&D Systems, MAB050), for Gαi with mouse monoclonal anti-Gαi (1:500) (Santa Cruz Biotechnology, sc-136478) or mono-ADP-ribose with a rabbit polyclonal anti-mono-ADP-ribose (1:1000) antibody (Hottiger-laboratory). Primary antibody membranes were washed thrice with TBST containing 2% (w/v) BSA for ten minutes each time. Primary antibody membranes were incubated with mouse IgG kappa binding protein conjugated to horseradish peroxidase (1:2500) (sc-516102, Santa Cruz Biotechnology) or goat anti-rabbit IgG conjugated to horseradish peroxidase (1:2500) (sc-2004, Santa Cruz Biotechnology) for 3 h at 4°C in rotation and washed thrice with TBST for ten minutes each time. All membranes were subsequently developed with WesternBright ECL (Advansta) and imaged on ImageQuant LAS 4000 (GE Healthcare). ***ii) substrate-excess condition with Western blot read-out*** Reactions (typically in 50 µL) contained 50 nM rPtxS1 proteins and 500 nM rGαi in 50 mM sodium phosphate (pH 7.0) and 1 μM biotinylated NAD^+^ (Trevigen, 4670-500-01). The reactions were carried at room temperature for 40 minutes. For validating hits from chemical screening, 10 μM of the indicated compound and 0.1% DMSO (control reaction) were used. Reactions were stopped by addition of Laemmli loading dye to 1 × and heating for 3 minutes at 90°C. The samples were run on SDS-PAGE and transferred to nitrocellulose membranes, followed by blocking with 1% (w/v) casein blocking buffer (Bio-Rad, 161-0782). Membranes were incubated with streptavidin conjugated to horse radish peroxidase (1:7000) (PerkinElmer, NEL750001EA) in 1% (w/v) casein blocking buffer (Bio-Rad, 161-0782) for 3 h at 4°C in rotation and washed thrice with TBST for ten minutes each time. Alternatively, for anti-His blotting, Penta His HRP conjugate (Qiagen, 34460) was diluted 1:5000 in 1% (w/v) casein blocking buffer (Bio-Rad, 161-0782). Blots were incubated at room temperature for 2 h, followed by 3 washes with TBST each for 5 minutes. Membranes were subsequently developed with WesternBright ECL (Advansta) and imaged on ChemiDoc XRS+ (Bio-Rad). ***iii) substrate-excess condition with nickel-plate readout*** The enzymatic activity of rPtxS1 in the absence (automodification) or presence of rGαi (substrate protein modification) and inhibitors (NSC288155 and NSC29193) was analyzed using 500 nM rPtxS1, 2 µM rGαi and 200 µM of inhibitors in 50 mM HEPES pH 7.5, 100 mM NaCl, 4 mM MgCl_2_ and 0.2 mM TCEP. Triplicate reactions were started at RT in PCR tubes by addition of an NAD^+^-mixture resulting to final concentrations of 24.5 µM NAD^+^ (Sigma, N3014) and 0.5 µM of biotinylated NAD^+^ (Trevigen, 4670-500-01). Reactions were incubated in constant shaking for 30 min at RT prior to addition to white 96-well nickel-coated plates (Pierce, 15242). Reactions were incubated on the nickel-coated plates for 60 min at RT after which reactions were stopped by addition of 7 M guanidine hydrochloride. Plates were washed after 5 min incubation at RT three times with TBST after which blocking solution containing 1% (w/v) BSA in TBST was added for 40 min. After blocking with BSA, streptavidin-conjugated horseradish peroxidase (GE Healthcare, RPN1231VS) in TBST-BSA was added (1:5000) for 60 min. Plates were washed four times with TBST after which chemiluminescent substrate (Advansta, WesternBright Quantum) was added and subsequent chemiluminescence was detected using Hidex Sense microplate reader (Hidex).

### *In vitro* NAD^+^ consumption assay

***i) basic reaction set-up*** Reactions were carried out in a U-shaped 96-well black plate (Greiner BioOne, 650209). Typically, reactions were conducted with 50 mM sodium phosphate (pH 7.0) at 25°C with shaking at 300 rpm with a reaction volume of 50 μL. The reactions were stopped by adding 20 μL of 20% acetophenone (diluted with ethanol) and 20 μL of 2M KOH and incubated at room temperature for 10 minutes. 90 μL of formic acid was added and further incubated for 20 minutes. The plates were read using Tecan infinity M1000 pro with excitation and emission wavelengths set at 372 and 444 nm, respectively. Maximum signal was defined as the NAD^+^ buffer control and the minimum signal was the rPtxS1-catalyzed reaction in the presence of rGαi. The raw fluorescence values were always subtracted from blank containing buffer. The assay was optimized to reach 60% NAD^+^ consumption. The optimized conditions for the assay are 125 nM rPtxS1, 500 nM NAD^+^ and 1 µM rGαi. The reaction has incubation time of 40 minutes at 25°C with shaking at 300 rpm. The DMSO concentration used was 0.1%. DMSO tolerance test was done with optimized conditions (0.1-5%). For rGαI substrate-independent NAD^+^ consumption activity of rPtxS1, identical conditions were used except that rGαi was excluded with longer incubation time of 60 minutes. Statistical analyses on typically 8 parallel values was conducted using two-tailed Student’s t-test two sample equal variance. ***ii)* validation of the NAD**^**+**^ **consumption assay** In order to establish repeatability of values for maximum and minimum signals between plates, wells and days, five control plates were tested. Experimental conditions and protein batches used for assay validation were the same. Three plates were made on one day while second and third days had one plate each. Each plate had 40 wells for maximum and minimum signals individually with buffer blanks. The CV% were calculated separately for minimum and maximum signals using (Standard deviation/average) *100. Assay parameters such as signal-to-noise (S/N), (S/B) signal-to-background, screening window coefficient (Z’) were calculated as described previously ^22, 23^. ***iii) small molecule library screening*** Each plate had buffer blank, control reaction (minimum signal, with rPtxS1-wt) and maximum signal (NAD^+^ and rGαi). Both maximum and minimum signals contain 0.1% DMSO. To correct for inherent fluorescence of compounds separate controls with NAD^+^, Gαi and compounds were prepared. Compounds were tested at a concentration of 10 μM. Compounds displaying inhibition more than 50% were considered as hits. Compound library for screening was obtained from the National Cancer Institute (NCI) Developmental Therapeutics program repository (https://dtp.cancer.gov). After the screening, we re-ordered the primary hit compounds in a powder form for subsequent *in vitro* and *in vivo* analyses. The hit compounds were analyzed to identify pan-assay interference compounds or aggregators (zinc15.docking.org). ***iv) IC***_***50***_ ***measurements*** Assay incubation time was adjusted such that the substrate conversion did not exceed 30% in order to minimize the effect of reduction in substrate concentration while maintaining a robust signal. Compounds were tested from a concentration range of (100-0.01 μM) in half-log dilution series. Each plate had buffer blank, positive control where PtxS1-wt is added which corresponds to 100% activity and negative control (without enzyme) where the activity was 0%. Control values were included as two half-log units below and above the inhibitor concentration series for reactions with 0% and 100% activity. IC_50_ values were obtained by fitting data to log (inhibitor) vs response - variable slope using GraphPad (GraphPad software, Inc.) Chemical structures were drawn using MarvinSketch 5.11.3 (chemAxon) or ChemDraw 18.0 (Perkin-Elmer).

### Structure preparation and molecular docking

The 2.7 Å resolution crystal structure of pertussis toxin was accessed from the Protein Data Bank (PDB) (PDB_1BCP ^15^). Herein, we considered only the catalytic PtxS1 subunit (1BCP, chain A). Residues 222-269 of PtxS1 were also removed from the coordinate file (see Fig. 1 and 2). The truncated structure was then prepared for docking using the protein preparation wizard ^38^ in Maestro (Schrödinger Release 2019-1: Maestro, Schrödinger, LLC, New York, NY, 2019). Hydrogen atoms were added, protonation states of ionizable groups were determined and the structure was energy minimized. A grid outlining the binding pocket was specified based on the superimposition of the PtxS1 structure with the S1 subunit of the pertussis-like toxin structure from *E. coli* with bound NAD^+^ (PDB_4Z9D; ^21^). The two toxins are 30% identical in primary amino acid sequence. The ligands NSC228155 (7-nitro-4-(1-oxidopyridin-1-ium-2-yl)sulfanyl-2,1,3-benzoxadiazole) and NSC29193 (purine-2,8-dithiol) were prepared for docking using the LigPrep program in Maestro (Schrödinger Release 2019-1: LigPrep, Schrödinger, LLC, New York, NY, 2019). Possible ionization states of the ligands at pH 7.0 ± 2.0 were determined and, in the case of NSC29193, four tautomeric states were also produced. The ligands were docked to the pertussis S1 structure using the glide XP and SP methods ^39^, keeping the protein structure rigid but allowing ligand flexibility, producing up to 20 poses for each ligand and tautomer. Since the resulting XP docking poses exhibited few interactions, the SP docking results were pursued further. The poses were ranked according to the glide docking score and free energy of binding calculations made with the Prime MM-GBSA module (Schrödinger Release 2019-1: Prime, Schrödinger, LLC, New York, NY, 2019).

### Molecular dynamics simulation

Molecular dynamics simulation (MDS) of the selected binding poses of NSC228155 and NSC29193 bound to the S1 structure was used to study the dynamics of the protein-ligand complexes using the Desmond program ^40^ in Maestro. The complexes were solvated using a TIP3P water model ^41^ in an octahedral box, with a 10 Å distance between solute surface atoms and an edge of the box. The systems were neutralized by adding Na^+^ counterions. Additional Na^+^/Cl^-^ ions were added to bring the systems to a 150 mM salt concentration. The simulations were carried out using the OPLS3e force field ^42^ at constant temperature (300 K) and constant pressure (1 atm), which were respectively regulated using the Nose-Hoover chain thermostat ^43^ and Martyna-Tobias-Klein barostat ^44^. Short-range and long-range interactions were computed with a 9 Å distance cutoff. The RESPA integrator ^45^ was employed with a 6.0 fs time step for long-range non-bonded interactions and a 2.0 fs time step for bonded and short range non-bonded interactions. The systems were relaxed with the default equilibration protocol in Desmond, followed by a 100 ns production simulation. Energies were saved every 1.2 ps, whereas coordinates were recorded every 100 ps. The resulting trajectories were analyzed in terms of ligand stability (as root mean squared deviation, RMSD) and lifetime of protein-ligand interactions using the Simulation Interactions Diagram application in Desmond.

### *In vivo* ADP-ribosylation assay

HEK293T cells grown in DMEM + 10% FBS were seeded in 3 mL volumes in 6-well plates (500 000 cells/well) in the late afternoon. The next morning fresh media containing NSC228155 at concentrations of 0.1, 1 and 5 μM was exchanged to the cells (0.05% DMSO in all reactions, including control reactions). Cells were incubated for 30 minutes at 37°C under normal cell culturing conditions, after which 10 ng/mL of pertussis AB_5_ holotoxin (List Biological Laboratories Inc., 179A) was added to the cells and incubation was continued for 2 h. Cells were then transferred on ice and washed twice with 1 × PBS and collected into 70 µL of lysis buffer [50 mM Tris-HCl (pH 7.5), 400 mM NaCl, 0.1% sodium deoxycholate, 1% NP-40, 75 μM tannic acid (PARG inhibitor), 40 μM PJ34 (PARP inhibitor) supplemented with Pierce Protease and Phosphatase Inhibitor Mini Tablets (40 μL/mL of stock solution – one tablet / 2 mL H_2_O, Thermo Scientific, 88668). Samples were kept on ice for 30 minutes and centrifuged for 15 minutes, 4°C, 16100 × g. Protein concentration was measured from the supernatants with Bradford protein assay. The samples were scaled for protein content to allow loading of 30 µg of total protein per lane in the SDS-PAGE. Laemmli loading dye was added to 1 × and the samples were boiled for 10 minutes at 95°C. The samples were run on SDS-PAGE and transferred to nitrocellulose membranes, followed by blocking with 4% (w/v) BSA in TBST. Membranes were probed in TBST containing 2% (w/v) BSA for mono-ADP-ribose with custom in-house rabbit polyclonal anti-mono-ADP-ribose (1:1000) (Hottiger-laboratory), for GAPDH with mouse monoclonal anti-GAPDH (1:1000) (Abcam, ab9484), for Gαi with mouse monoclonal anti-Gαi (1:500) (Santa Cruz Biotechnology, sc-136478), for poly-ADP-ribose with rabbit polyclonal anti-PAR (1:1000) (Enzo Life Sciences, ALX-210-890A) and for PARP1 with mouse monoclonal anti-PARP1 (1:300) (Santa Cruz Biotechnology, sc-8007). Membranes were washed thrice with TBST containing 2% (w/v) BSA for ten minutes each time. Membranes were incubated in TBST containing 2% (w/v) BSA with mouse IgG kappa binding protein conjugated to horseradish peroxidase (1:5000) (sc-516102, Santa Cruz Biotechnology) or goat anti-rabbit IgG conjugated to horseradish peroxidase (1:5000) (sc-2004, Santa Cruz Biotechnology) for 3 h at 4°C on a rotary and washed thrice with TBST for ten minutes each time. Membranes were subsequently developed with WesternBright ECL (Advansta) and imaged on ImageQuant LAS 4000 (GE Healthcare). Pixel intensities were quantified form the Western blot TIFF-files using ImageJ 1.44o (NIH, USA, https://imagej.nih.gov/ij/index.html).

### MTT cell viability assay

HEK293T cells grown in DMEM + 10% FBS were seeded in 100 μL volumes in 96-well plates (20 000 cells/well) in the late afternoon. The next morning fresh media containing varying concentrations (0, 0.1, 0.5, 1, 2.5, 5, 10, 20, 40, 60 μM) of NSC228155 was added in triplicate. In all the wells final concentration of DMSO was 0.6%. Cells were incubated for 2.5 h at 37°C under normal cell culturing conditions. Cell viability was investigated using the CellTiter 96 Non-Radioactive Cell Proliferation Assay (MTT) (Promega, G4002) according to the manufacturer’s instructions. Incubation time with the MTT dye solution was 1Uh at 37°C under normal cell culturing conditions. NSC228155 cytotoxicity IC_50_ value was calculated using GraphPad (GraphPad software, Inc.) Data are displayed as means and the standard error of the mean. Data were normalized to cells treated with 0.6% DMSO (negative controls, 100% viability) and cells killed with 200 μM H_2_O_2_ (positive control, 0% viability) and expressed as percentage of these controls. Normalized response is compared to common log of the inhibitor concentration (µM) and the IC_50_ calculated using a variable slope. Statistical analyses were conducted using two-tailed Student’s t-test two sample equal variance.

## ANCILLARY INFORMATION

### Description of the Supporting Information

Figure S1. SEC-analysis of rPtxS1-rGαi complex formation in solution.

Figure S2. Catalytic activity of rPtxS1 is dependent on two acidic amino acids.

Figure S3. rPtxS1 is incapable of ADP-ribosylating a C351A-mutant of rGαi.

Figure S4. Molecular dynamics simulation data.

Table S1. Assay performance statistic of the *in vitro* NAD^+^ consumption assay.

Table S2. ARTD/PARP inhibitory compounds analyzed for rPtxS1 inhibition.

## AUTHOR CONTRIBUTIONS

Yashwanth Ashok and Moona Miettinen contributed equally to this study.

## ACKNOWLEDGEMENTS

The research in the laboratory of A.T.P. is financially supported by Academy of Finland (grant no. 295296), Sigrid Juselius Foundation, Turku Doctoral Programme of Molecular Medicine (TuDMM) (to M.M.) and University of Turku, Turku, Finland. This work was funded in the L.L. laboratory by Academy of Finland (grant no. 287063, 294085 and 319299). The use of the facilities of the Biocenter Oulu for DNA sequencing, Proteomics and Protein Analysis and Protein Crystallography, a member of Biocenter Finland and Instruct-FI, are gratefully acknowledged. The laboratory of M.S.J. is supported by Sigrid Juselius Foundation, Joe, Pentti and Tor Memorial Fund, and Doctoral Network of Informational and Structural Biology (to M.T., Åbo Akademi Graduate School); computational infrastructure and core faculty support from Biocenter Finland (bioinformatics, structural biology and drug discovery and chemical biology nodes), CSC IT Center for Science; Academy of Finland FIRI infrastructure funding (grant no. 320005); screening core faculty of Biocity Turku, and Drug Discovery and Diagnostics strategic funding to Åbo Akademi University. We thank Dr. Jukka Lehtonen for his scientific IT support. The funders had no role in study design, data collection and interpretation, or the decision to submit the work for publication. The authors declare that they have no conflicts of interest.

## ABBREVIATIONS

ART: ADP-ribosyltransferase
ADP: adenosine diphosphate
*B. pertussis*: Bordetella pertussis
BSA: bovine serum albumin
DMSO: dimethyl sulfoxide
DSF: differential scanning fluorimetry
*E. coli*: Escherichia coli
Gαi: inhibitory α-subunit of heterotrimeric G-protein
GPCR: G-protein coupled receptor
HRP: horse radish peroxidase
IC_50_: half maximal inhibitory concentration
LB: Luria-Bertani
MALS: multi-angle light scattering
MAR: mono-ADP-ribose
MDS: molecular dynamic simulation
NAD^+^: nicotinamide adenine dinucleotide
PAR: poly-ADP-ribose
PARP1: poly(ADP-ribose) polymerase 1 (also known as ARTD1)
PCR: polymerase chain reaction
PDB: protein data bank
PtxS1: pertussis toxin S1 subunit
rGαi: recombinant inhibitory α-subunit of heterotrimeric G-protein
RMSD: root-mean-square deviation
rPtxS1: recombinant pertussis toxin S1 subunit
SDS-PAGE: sodium dodecyl sulphate polyacrylamide gel electrophoresis
SEC: size exclusion chromatography
TBST: Tris-buffered saline containing 0.05 % Tween
Tm: melting temperature

## REFERENCES

1. Kilgore, P. E.; Salim, A. M.; Zervos, M. J.; Schmitt, H. J., Pertussis: Microbiology, disease, treatment, and prevention. Clin Microbiol Rev 2016, 29 (3), 449–86.

2. Yeung, K. H. T.; Duclos, P.; Nelson, E. A. S.; Hutubessy, R. C. W., An update of the global burden of pertussis in children younger than 5 years: a modelling study. Lancet Infect Dis 2017, 17 (9), 974–80.

3. Skoff, T. H.; Baumbach, J.; Cieslak, P. R., Tracking pertussis and evaluating control measures through enhanced pertussis surveillance, emerging infections program, United States. Emerg Infect Dis 2015, 21 (9), 1568–73.

4. Guillot, S.; Descours, G.; Gillet, Y.; Etienne, J.; Floret, D.; Guiso, N., Macrolide-resistant *Bordetella pertussis* infection in newborn girl, France. Emerg Infect Dis 2012, 18 (6), 966–8.

5. Wang, Z.; Li, Y.; Hou, T.; Liu, X.; Liu, Y.; Yu, T.; Chen, Z.; Gao, Y.; Li, H.; He, Q., Appearance of macrolide-resistant *Bordetella pertussis* strains in China. Antimicrob Agents Chemother 2013, 57 (10), 5193–4.

6. Scanlon, K.; Skerry, C.; Carbonetti, N., Association of pertussis toxin with severe pertussis disease. Toxins (Basel) 2019, 11 (7).

7. Bouchez, V.; Brun, D.; Cantinelli, T.; Dore, G.; Njamkepo, E.; Guiso, N., First report and detailed characterization of *B. pertussis* isolates not expressing pertussis toxin or pertactin. Vaccine 2009, 27 (43), 6034–41.

8. Morse, S. I.; Morse, J. H., Isolation and properties of the leukocytosis- and lymphocytosis-promoting factor of *Bordetella pertussis*. J Exp Med 1976, 143 (6), 1483–502.

9. Hall, E.; Parton, R.; Wardlaw, A. C., Cough production, leucocytosis and serology of rats infected intrabronchially with *Bordetella pertussis*. J Med Microbiol 1994, 40 (3), 205–13.

10. Parton, R.; Hall, E.; Wardlaw, A. C., Responses to *Bordetella pertussis* mutant strains and to vaccination in the coughing rat model of pertussis. J Med Microbiol 1994, 40 (5), 307–12.

11. Scanlon, K. M.; Snyder, Y. G.; Skerry, C.; Carbonetti, N. H., Fatal pertussis in the neonatal mouse model is associated with pertussis toxin-mediated pathology beyond the airways. Infect Immun 2017, 85 (11).

12. Stein, P. E.; Boodhoo, A.; Armstrong, G. D.; Cockle, S. A.; Klein, M. H.; Read, R. J., The crystal structure of pertussis toxin. Structure 1994, 2 (1), 45–57.

13. Stein, P. E.; Boodhoo, A.; Armstrong, G. D.; Heerze, L. D.; Cockle, S. A.; Klein, M. H.; Read, R. J., Structure of a pertussis toxin-sugar complex as a model for receptor binding. Nat Struct Biol 1994, 1 (9), 591–6.

14. Weiss, A. A.; Johnson, F. D.; Burns, D. L., Molecular characterization of an operon required for pertussis toxin secretion. Proc Natl Acad Sci U S A 1993, 90 (7), 2970–4.

15. Hazes, B.; Boodhoo, A.; Cockle, S. A.; Read, R. J., Crystal structure of the pertussis toxin-ATP complex: a molecular sensor. J Mol Biol 1996, 258 (4), 661–71.

16. Simon, N. C.; Aktories, K.; Barbieri, J. T., Novel bacterial ADP-ribosylating toxins: structure and function. Nat Rev Microbiol 2014, 12 (9), 599–611.

17. Katada, T.; Ui, M., Direct modification of the membrane adenylate cyclase system by islet-activating protein due to ADP-ribosylation of a membrane protein. Proc Natl Acad Sci U S A 1982, 79 (10), 3129–33.

18. West, R. E.; Moss, J.; Vaughan, M.; Liu, T.; Liu, T. Y., Pertussis toxin-catalyzed ADP-ribosylation of transducin. Cysteine 347 is the ADP-ribose acceptor site. J Biol Chem 1985, 260 (27), 14428–30.

19. Graf, R.; Codina, J.; Birnbaumer, L., Peptide inhibitors of ADP-ribosylation by pertussis toxin are substrates with affinities comparable to those of the trimeric GTP-binding proteins. Mol Pharmacol 1992, 42 (5), 760–4.

20. Weis, W. I.; Kobilka, B. K., The molecular basis of G protein-coupled receptor activation. Annu Rev Biochem 2018, 87, 897–919.

21. Littler, D. R.; Ang, S. Y.; Moriel, D. G.; Kocan, M.; Kleifeld, O.; Johnson, M. D.; Tran, M. T.; Paton, A. W.; Paton, J. C.; Summers, R. J.; Schembri, M. A.; Rossjohn, J.; Beddoe, T., Structure-function analyses of a pertussis-like toxin from pathogenic *Escherichia coli* reveal a distinct mechanism of inhibition of trimeric G-proteins. J Biol Chem 2017, 292 (36), 15143–58.

22. Putt, K. S.; Hergenrother, P. J., An enzymatic assay for poly(ADP-ribose) polymerase-1 (PARP-1) via the chemical quantitation of NAD(+): application to the high-throughput screening of small molecules as potential inhibitors. Anal Biochem 2004, 326 (1), 78–86.

23. Venkannagari, H.; Fallarero, A.; Feijs, K. L.; Lüscher, B.; Lehtiö, L., Activity-based assay for human mono-ADP-ribosyltransferases ARTD7/PARP15 and ARTD10/PARP10 aimed at screening and profiling inhibitors. Eur J Pharm Sci 2013, 49 (2), 148–56.

24. Pizza, M.; Covacci, A.; Bartoloni, A.; Perugini, M.; Nencioni, L.; De Magistris, M. T.; Villa, L.; Nucci, D.; Manetti, R.; Bugnoli, M., Mutants of pertussis toxin suitable for vaccine development. Science 1989, 246 (4929), 497–500.

25. Locht, C.; Keith, J. M., Pertussis toxin gene: nucleotide sequence and genetic organization. Science 1986, 232 (4755), 1258–64.

26. Nicosia, A.; Perugini, M.; Franzini, C.; Casagli, M. C.; Borri, M. G.; Antoni, G.; Almoni, M.; Neri, P.; Ratti, G.; Rappuoli, R., Cloning and sequencing of the pertussis toxin genes: operon structure and gene duplication. Proc Natl Acad Sci U S A 1986, 83 (13), 4631–5.

27. Moss, J.; Stanley, S. J.; Burns, D. L.; Hsia, J. A.; Yost, D. A.; Myers, G. A.; Hewlett, E. L., Activation by thiol of the latent NAD glycohydrolase and ADP-ribosyltransferase activities of *Bordetella* pertussis toxin (islet-activating protein). J Biol Chem 1983, 258 (19), 11879–82.

28. Finck-Barbançon, V.; Barbieri, J. T., Preferential processing of the S1 subunit of pertussis toxin that is bound to eukaryotic cells. Mol Microbiol 1996, 22 (1), 87–95.

29. Sakanyan, V.; Hulin, P.; Alves de Sousa, R.; Silva, V. A.; Hambardzumyan, A.; Nedellec, S.; Tomasoni, C.; Logé, C.; Pineau, C.; Roussakis, C.; Fleury, F.; Artaud, I., Activation of EGFR by small compounds through coupling the generation of hydrogen peroxide to stable dimerization of Cu/Zn SOD1. Sci Rep 2016, 6, 21088.

30. Lüscher, B.; Bütepage, M.; Eckei, L.; Krieg, S.; Verheugd, P.; Shilton, B. H., ADP-ribosylation, a multifaceted posttranslational modification involved in the control of cell physiology in health and disease. Chem Rev 2018, 118 (3), 1092–1136.

31. Winter, K.; Zipprich, J.; Harriman, K.; Murray, E. L.; Gornbein, J.; Hammer, S. J.; Yeganeh, N.; Adachi, K.; Cherry, J. D., Risk factors associated with infant deaths from pertussis: a case-control study. Clin Infect Dis 2015, 61 (7), 1099–106.

32. Acquaye-Seedah, E.; Huang, Y.; Sutherland, J. N.; DiVenere, A. M.; Maynard, J. A., Humanised monoclonal antibodies neutralise pertussis toxin by receptor blockade and reduced retrograde trafficking. Cell Microbiol 2018, 20 (12), e12948.

33. Nguyen, A. W.; Wagner, E. K.; Laber, J. R.; Goodfield, L. L.; Smallridge, W. E.; Harvill, E. T.; Papin, J. F.; Wolf, R. F.; Padlan, E. A.; Bristol, A.; Kaleko, M.; Maynard, J. A., A cocktail of humanized anti-pertussis toxin antibodies limits disease in murine and baboon models of whooping cough. Sci Transl Med 2015, 7 (316), 316ra195.

34. Turgeon, Z.; Jørgensen, R.; Visschedyk, D.; Edwards, P. R.; Legree, S.; McGregor, C.; Fieldhouse, R. J.; Mangroo, D.; Schapira, M.; Merrill, A. R., Newly discovered and characterized antivirulence compounds inhibit bacterial mono-ADP-ribosyltransferase toxins. Antimicrob Agents Chemother 2011, 55 (3), 983–91.

35. Pinto, A. F.; Ebrahimi, M.; Saleeb, M.; Forsberg, Å.; Elofsson, M.; Schüler, H., Identification of inhibitors of *Pseudomonas aeruginosa* exotoxin-S ADP-ribosyltransferase activity. J Biomol Screen 2016, 21 (6), 590–5.

36. Locht, C.; Cieplak, W.; Marchitto, K. S.; Sato, H.; Keith, J. M., Activities of complete and truncated forms of pertussis toxin subunits S1 and S2 synthesized by *Escherichia coli*. Infect Immun 1987, 55 (11), 2546–53.

37. Pulliainen, A. T.; Pieles, K.; Brand, C. S.; Hauert, B.; Böhm, A.; Quebatte, M.; Wepf, A.; Gstaiger, M.; Aebersold, R.; Dessauer, C. W.; Dehio, C., Bacterial effector binds host cell adenylyl cyclase to potentiate Gαs-dependent cAMP production. Proc Natl Acad Sci U S A 2012, 109 (24), 9581–6.

38. Sastry, G. M.; Adzhigirey, M.; Day, T.; Annabhimoju, R.; Sherman, W., Protein and ligand preparation: parameters, protocols, and influence on virtual screening enrichments. J Comput Aided Mol Des 2013, 27 (3), 221–34.

39. Friesner, R. A.; Banks, J. L.; Murphy, R. B.; Halgren, T. A.; Klicic, J. J.; Mainz, D. T.; Repasky, M. P.; Knoll, E. H.; Shelley, M.; Perry, J. K.; Shaw, D. E.; Francis, P.; Shenkin, P. S., Glide: a new approach for rapid, accurate docking and scoring. 1. Method and assessment of docking accuracy. J Med Chem 2004, 47 (7), 1739–49.

40. Bowers KJ, Chow E, Xu H, Dror RO, Eastwood MP, Gregersen BA, Klepeis JL, Kolossvary I, Moraes MA, Sacerdoti FD, Salmon JoK, Shan Y, Shaw DE. Scalable algorithms for molecular dynamics simulations on commodity clusters. Proceedings of the ACM/IEEE Conference on Supercomputing (SC06), Tampa, Florida, 2006, November 11-17

41. Jorgensen, W. L.; Chandrasekhar, J.; Madura, J. D.; Impey, R. W.; Klein, M., Comparison of simple potential functions for simulating liquid water. J Chem Phys 1983, 79: 926–35.

42. Harder, E.; Damm, W.; Maple, J.; Wu, C.; Reboul, M.; Xiang, J. Y.; Wang, L.; Lupyan, D.; Dahlgren, M. K.; Knight, J. L.; Kaus, J. W.; Cerutti, D. S.; Krilov, G.; Jorgensen, W. L.; Abel, R.; Friesner, R. A., OPLS3: A force field providing broad coverage of drug-like small molecules and proteins. J Chem Theory Comput 2016, 12 (1), 281–96.

43. Martyna, G. J.; Klein, M. L.; Tuckerman, M., Nose-Hoover chains - the canonical ensemble via continuous dynamics. J Chem Phys 1992, 97: 2635–43.

44. Martyna, G. J.; Tobias, D. J.; Klein, M. L., Constant-pressure molecular dynamics algorithms. J Chem Phys 1994, 101: 4177–89.

45. Tuckerman, M.; Berne, B. J.; Martyna, G. J., Reversible multiple time scale molecular dynamics. J Chem Phys 1992, 97:1990–2001.

