## Supporting Information for "Discovery of compounds inhibiting the ADP-ribosyltransferase activity of pertussis toxin"

**Supporting Information** – Figures S1-S4, Table S1-S2

Figure S1. SEC-analysis of rPtxS1-rGαi complex formation in solution.

Figure S2. Catalytic activity of rPtxS1 is dependent on two acidic amino acids.

Figure S3. rPtxS1 is incapable of ADP-ribosylating a C351A-mutant of rGαi.

Figure S4. Molecular dynamics simulation data.

Table S1. Assay performance statistic of the *in vitro* $NAD^+$ consumption assay.

Table S2. ARTD/PARP inhibitory compounds analyzed for rPtxS1 inhibition.

### Discovery of compounds inhibiting the ADP-ribosyltransferase activity of pertussis toxin

Yashwanth Ashok[a,#], Moona Miettinen[b,c,#], Danilo Kimio Hirabae de Oliveira[a], Mahlet Z. Tamirat[d], Katja Näreoja[b], Avlokita Tiwari[b], Michael O. Hottiger[e], Mark S. Johnson[d], Lari Lehtiö[a,*] & Arto T. Pulliainen[b,*]

[a]Faculty of Biochemistry and Molecular Medicine, Biocenter Oulu, University of Oulu, Oulu, Finland

[b]Institute of Biomedicine, Research Center for Cancer, Infections, and Immunity, University of Turku, Turku, Finland

[c]Turku Doctoral Programme of Molecular Medicine (TuDMM), University of Turku, Turku, Finland

[d]Structural Bioinformatics Laboratory, Biochemistry, Faculty of Science and Engineering, Åbo Akademi University, Turku, Finland

[e]Department of Molecular Mechanisms of Disease, University of Zurich, Zurich, Switzerland

[#] Equal contribution, [*] Corresponding author

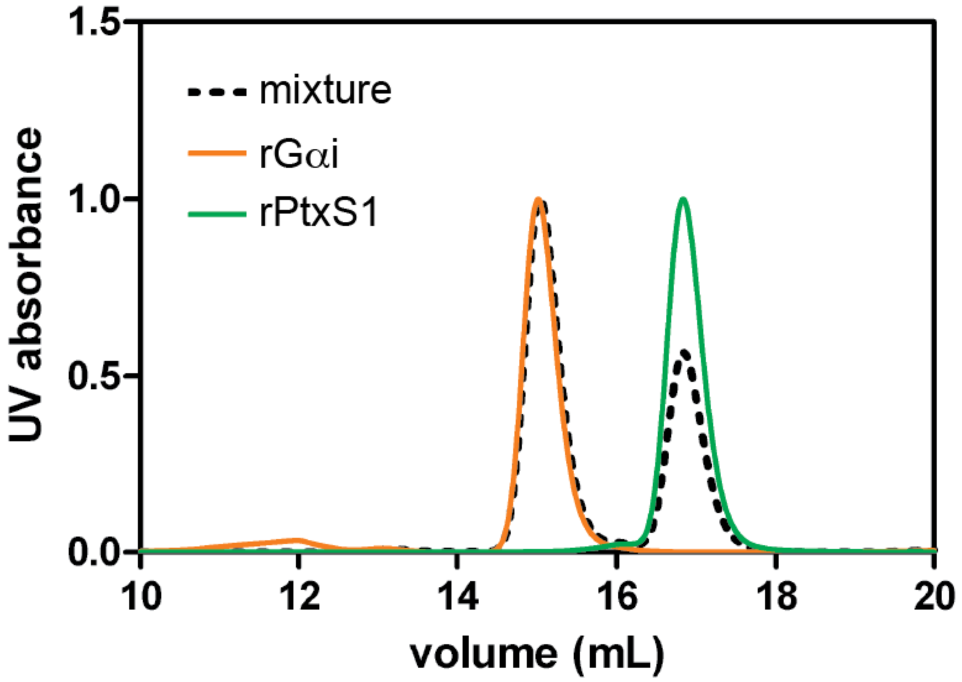


**Figure S1. SEC-analysis of rPtxS1-rGαi complex formation in solution.** Proteins were injected into the column either alone (100 µg) or in a mixture (100 µg + 100 µg).

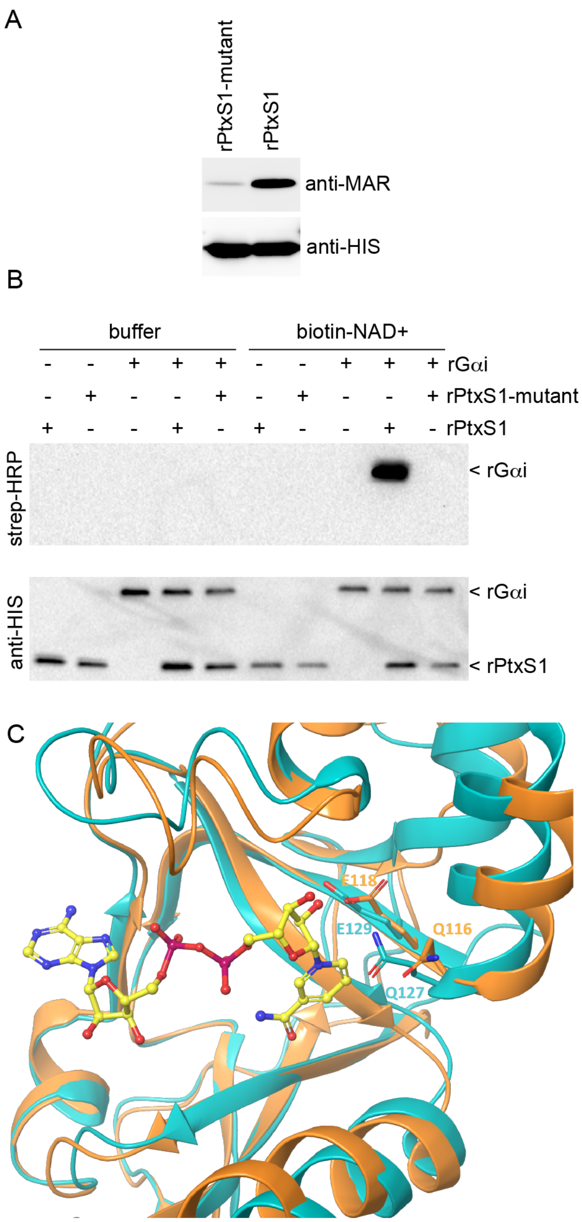


**Figure S2. Catalytic activity of rPtxS1 is dependent on two acidic amino acids. A)** Effect of Q127D/E129D mutation to auto-ADP-ribosylation of rPtxS1 that took place inside the *E. coli* expression host. **B)** Effect of Q127D/E129D mutation to rPtxS1-catalyzed ADP-ribosylation of rGαi *in vitro* (substrate-excess conditions). Blots probed, stripped and re-probed in the order of 1) strep-HRP and 2) anti-HIS. **C)** Structural comparison of the NAD$^+$-binding pocket of pertussis-like toxin from *E. coli* (PDB_4Z9C, PDB_4Z9D) with the corresponding area of PtxS1 (PDB_1BCP, cyan). NAD$^+$-binding pose from Q116/E118 mutant structure of pertussis-like toxin from *E. coli* (PDB_4Z9D) was superimposed on top of the wild-type structure (PDB_4Z9C, orange).

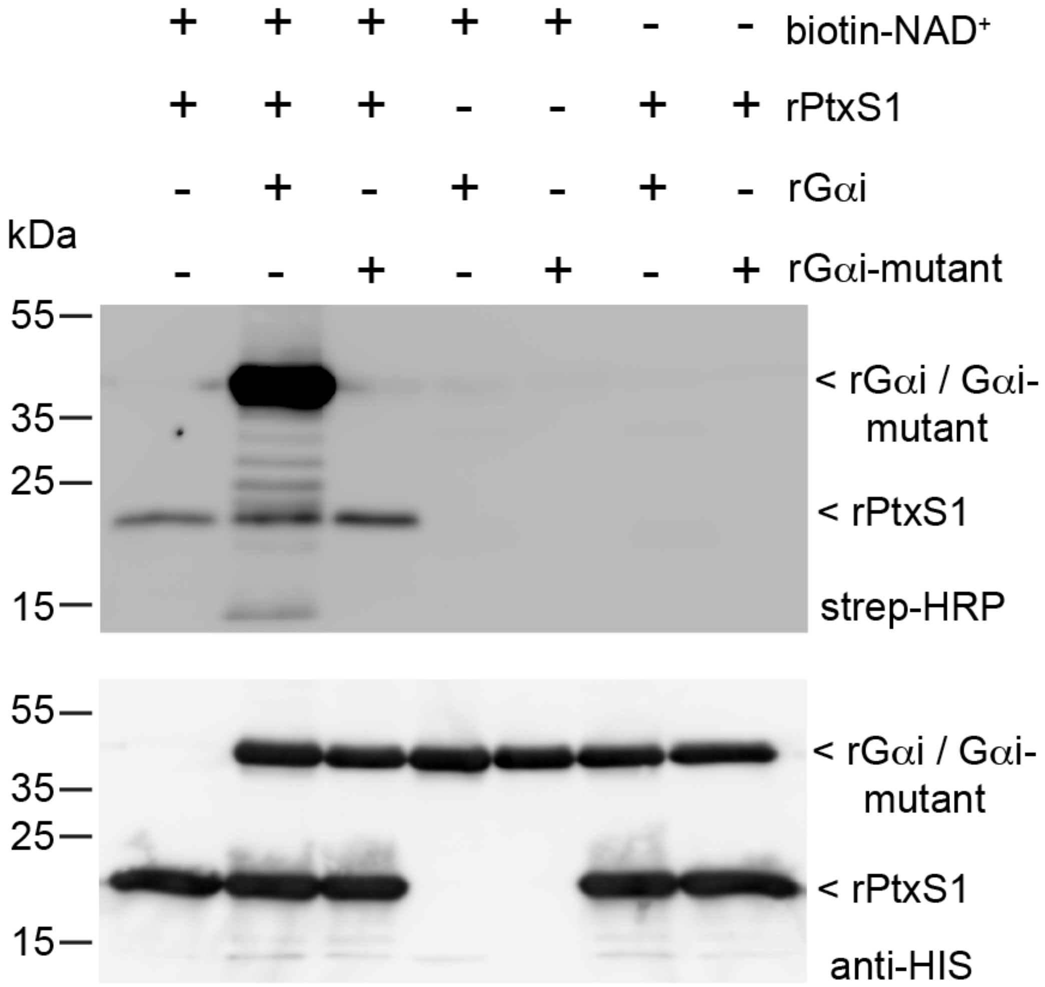


**Figure S3. rPtxS1 is incapable of ADP-ribosylating a C351A-mutant of rGαi.** Effect of C351A mutation to rPtxS1-catalyzed ADP-ribosylation of rGαi *in vitro* (enzyme-excess conditions). Blots probed, stripped and re-probed in the order of 1) strep-HRP and 2) anti-HIS.

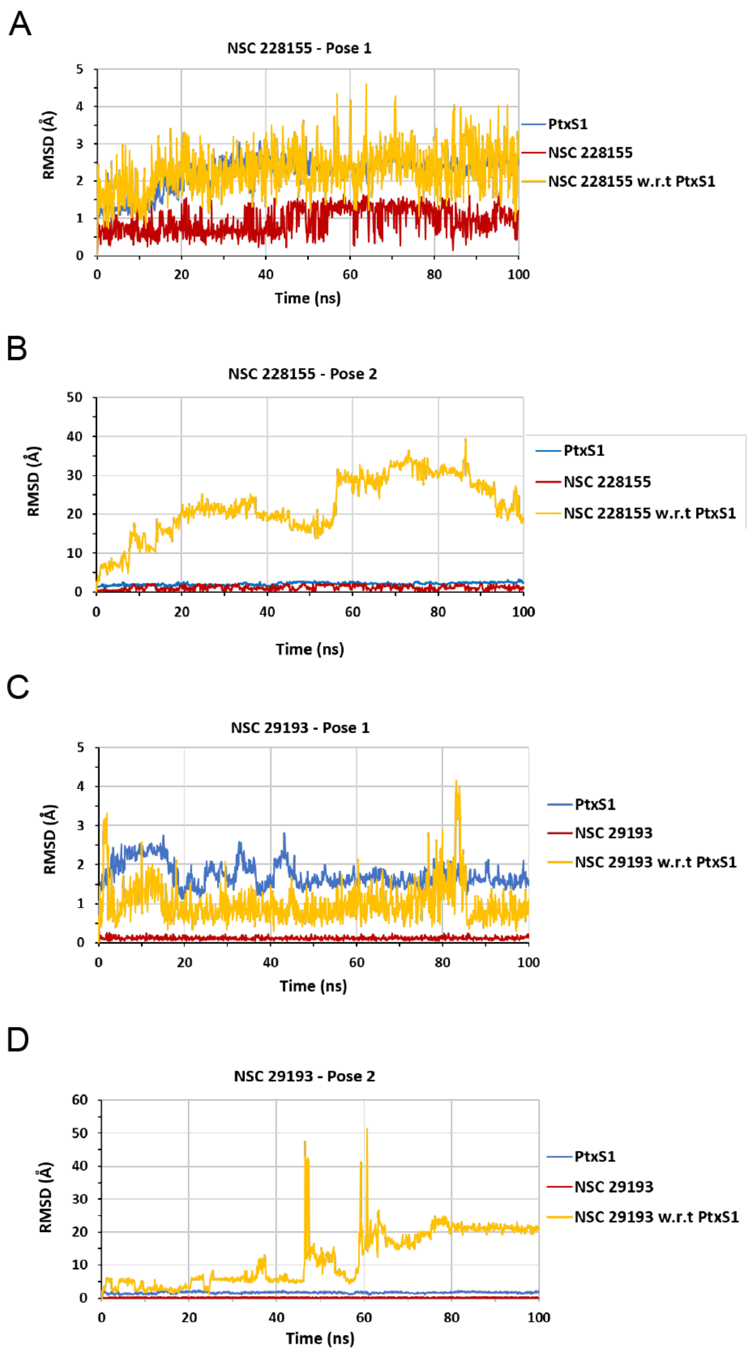


**Figure S4. Molecular dynamics simulation data.** Root mean square deviation (RMSD) of pose 1 **A)** and pose 2 **B)** of NSC228155 and pose 1 **C)** and pose 2 **D)** NSC29193 (see Fig. 5). Backbone atom RMSD of PtxS1 (blue) and the ligands (NSC228155 / NSC29193 - red) based on their first frame structures. Ligand RMSD (yellow) when superimposition is based on pertussis backbone atoms.

**Table S1. Assay performance statistic of the *in vitro* $NAD^+$ consumption assay.** In order to establish repeatability of values for maximum and minimum signals between plates, wells and days, five control plates were tested. Experimental conditions and protein batches used for assay validation were the same. Three plates were made on one day while second and third days had one plate each.

| **Parameter** | **Values** |
|---|---|
| S/B | 2.60 ± 0.2 |
| S/N | 13.17±1.5 |
| Z' | 0.68±0.0 |
| Well-to-well CV% max | 3.5±0.9 |
| Well-to-well CV% min | 7.8±0.9 |
| Plate-to-plate CV% * | 1.2 |
| Day-to-day CV% * | 5.6 (5.4 – 5.9)[#] |

*calculated from Z' values

[#]indicates range for Day-to-Day CV

#### Table S2. ARTD/PARP inhibitory compounds analyzed for rPtxS1 inhibition.

| Compound name (IUPAC) | Acronym | Supplier |
|---|---|---|
| 3-aminobenzamide | 3-AB | ALEXIS Biochemicals |
| Nicotinamide | | ALEXIS Biochemicals |
| Benzamide | | ALEXIS Biochemicals |
| 2-[(2R)-2-methylpyrrolidin-2-yl]-1H-1,3-benzodiazole-4-carboxamide | Veliparib | Medchemtronica |
| 5-amino-1,2-dihydroisoquinolin-1-one | 5-AIQ | ALEXIS Biochemicals |
| 5-amino-3-methyl-1,2-dihydroisoquinolin-1-one | 3-Methyl-5-AIQ | ALEXIS Biochemicals |
| 8-amino-3-azatricyclo[7.3.1.0^{5,13}]trideca-1(12),5,7,9(13),10-pentaene-2,4-dione | 4-ANI | ALEXIS Biochemicals |
| 5-[4-(piperidin-1-yl)butoxy]-1,2,3,4-tetrahydroisoquinolin-1-one | DPQ | ALEXIS Biochemicals |
| 2-methyl-1H,4H,5H,7H,8H-thiopyrano[4,3-d]pyrimidin-4-one | DR2313, DRL | ALEXIS Biochemicals |
| 2-(4-{[(2S,3S,4R,5R)-5-(6-amino-9H-purin-9-yl)-3,4-dihydroxyoxolan-2-yl]carbonyl}piperazin-1-yl)-N-(1-oxo-2,3-dihydro-1H-isoindol-4-yl)acetamide | EB-47 | ALEXIS Biochemicals |
| 1,4-dihydroquinazolin-4-one | 4-Hydroxyquinazoline | ALEXIS Biochemicals |
| 6-amino-5-iodo-2H-chromen-2-one | INH2BP | ALEXIS Biochemicals |
| 5-hydroxy-1,2-dihydroisoquinolin-1-one | 1,5-Isoquinolinediol | ALEXIS Biochemicals |
| (2E,4S,4aS,5aR,12aS)-2-[amino(hydroxy)methylidene]-4,7-bis(dimethylamino)-10,11,12a-trihydroxy-1,2,3,4,4a,5,5a,6,12,12a-decahydrotetracene-1,3,12-trione | Minocin | ALEXIS Biochemicals |
| 8-hydroxy-2-methyl-1,4-dihydroquinazolin-4-one | NU1025, 4PAX | ALEXIS Biochemicals |
| 5,6-dihydrophenanthridin-6-one | phenanthridone | ALEXIS Biochemicals |
| 2-(dimethylamino)-N-(6-oxo-5,6-dihydrophenanthridin-2-yl)acetamide | PJ-34, P34 | ALEXIS Biochemicals |
| 4H,5H-thieno[2,3-c]isoquinolin-5-one | TIQ-A | ALEXIS Biochemicals |
| 1,7-dimethyl-2,3,6,7-tetrahydro-1H-purine-2,6-dione | 1,7-dimethylxanthine | Sigma |
| 3-(4-chlorophenyl)quinoxaline-5-carboxamide | CNQ | Calbiochem / WVR |
| 4-({3-[(4-cyclopropanecarbonylpiperazin-1-yl)carbonyl]-4-fluorophenyl}methyl)-1,2-dihydrophthalazin-1-one | Olaparib | Medchemtronica |
| (4Z)-4-[(1-methyl-1H-pyrrol-2-yl)methylidene]-1,2,3,4-tetrahydroisoquinoline-1,3-dione | BYK204165 | Sigma |
| 2-[4-(trifluoromethyl)phenyl]-1H,4H,5H,7H,8H-thiopyrano[4,3-d]pyrimidin-4-one | XAV939 | Maybridge |
| 2-(pyridin-2-yl)-5H,7H,8H-thiopyrano[4,3-d]pyrimidin-4-ol | RF03877 | Maybridge |
| 2-cyclopropyl-5H,7H,8H-thiopyrano[4,3-d]pyrimidin-4-ol | RF03876 | Maybridge |
| 4-[(1R,2S,6R,7S)-3,5-dioxo-4-azatricyclo[5.2.1.0^{2,6}]dec-8-en-4-yl]-N-(quinolin-8-yl)benzamide | IWR-1 | Sigma |
| N-(6-methyl-1,3-benzothiazol-2-yl)-2-({4-oxo-3-phenyl-3H,4H,6H,7H-thieno[3,2-d]pyrimidin-2-yl}sulfanyl)acetamide | IWP-2 | Sigma |
| 4-iodo-3-nitrobenzamide | Iniparib | Selleck Biochemicals |
| 6-fluoro-2-{4-[(methylamino)methyl]phenyl}-3,10-diazatricyclo[6.4.1.0^{4,13}]trideca-1,4(13),5,7-tetraen-9-one | Rucaparib, AG014699 | Medchemtronica |
| 1-oxo-1,2-dihydroisoquinolin-5-yl benzoate | UPF1035 | Enzo Life Sciences |
| 5-(2-oxo-2-phenylethoxy)-1,2-dihydroisoquinolin-1-one | UPF1069 | Enzo Life Sciences |
| 2-phenyl-4H-chromen-4-one | Flavone | Sigma |
